# Chemoproteomic profiling reveals cellular targets of nitro-fatty acids

**DOI:** 10.1101/2021.07.12.451990

**Authors:** Mingyu Fang, Kuan Hsun Huang, Wei-Ju Tu, Yi-Ting Chen, Pei-Yun Pan, Wan-Chi Hsiao, Yi-Yu Ke, Lun K. Tsou, Mingzi M. Zhang

## Abstract

Nitro-fatty acids are a class of endogenous electrophilic lipid mediators with anti-inflammatory and cytoprotective effects in a wide range of inflammatory and fibrotic disease models. While these beneficial biological effects of nitro-fatty acids are mainly attributed to their ability to form covalent adducts with proteins, only a small number of proteins are known to be nitro-alkylated and the scope of protein nitro-alkylation remains undetermined. Here we describe the synthesis and application of a clickable nitro-fatty acid probe for the detection and first global identification of mammalian proteins that are susceptible to nitro-alkylation. 184 high confidence nitro-alkylated proteins were identified in human macrophages, majority of which are novel targets of nitro-fatty acids, including Extended synaptotagmin 2 (ESYT2), Signal transducer and activator of transcription 3 (STAT3), Toll-like receptor 2 (TLR2), Retinoid X receptor alpha (RXRα) and Glucocorticoid receptor (NR3C1). In particular, we showed that 9-nitro-oleate covalently modified and inhibited dexamethasone binding to NR3C1. Bioinformatic analyses revealed that nitro-alkylated proteins are highly enriched in endoplasmic reticulum and transmembrane proteins, and are overrepresented in lipid metabolism and transport pathways. This study significantly expands the scope of protein substrates targeted by nitro-fatty acids in living cells and provides a useful resource towards understanding the pleiotropic biological roles of nitro-fatty acids as signaling molecules or as multi-target therapeutic agents.

## 1. Introduction

Nitro-fatty acids, or nitroalkene fatty acids, are a class of electrophilic lipid mediators with potent anti-inflammatory and cytoprotective effects. Produced endogenously via non-enzymatic nitration of unsaturated lipids during digestion, cellular metabolism and inflammation, nitro-fatty acids have demonstrated beneficial bioactivities in a wide range of cardiovascular, pulmonary, renal and hepatic inflammatory and fibrotic disease models.^1-4^ In particular, 10-nitro-oleate (CXA-10) is being explored as a therapeutic against a number of inflammatory diseases in phase II clinical trials.^4^ While it is increasingly recognized that exogenous and endogenously formed nitro-fatty acids downregulate inflammatory signaling pathways to alleviate chronic inflammation, mechanistic insights into how these lipids exert their functions through pleiotropic targets in different tissues and cell types will inform the development of therapeutic strategies.

The formation of nitro-fatty acid-protein adducts, or nitro-alkylation, represents the primary mechanism by which nitroalkene fatty acids exert their biological functions (Supplementary Figure 1). Possessing strong electron-withdrawing nitro groups, nitro-fatty acids are robust Michael acceptors that react with nucleophilic amino acids, predominantly cysteines and sometimes histidines, with reaction rates exceeding those of hydrogen peroxide and other non-nitrated lipid-derived electrophiles such as 4-hydroxy-2-nonenal (HNE).^5^ A number of proteins have been reported to form covalent adducts with nitro-fatty acids, which resulted in changes to their stability, structure or function.^3,6^ Nevertheless, the overall scope of protein nitro-alkylation remains undetermined. Most studies focused on individual well-characterized protein targets, particularly Peroxisome proliferator-activated receptor gamma (PPARγ), Kelch-like ECH-associated protein 1 (KEAP1), Nuclear factor kappa B (NF-κB), and their associated signaling pathways to explain the interesting range of bioactivities observed for nitro-fatty acids. Yet there is increasing evidence that context-specific nitro-alkylation events can contribute to the effects of nitro-fatty acids on cellular function and host physiology. For example, despite its central role in modulating inflammatory responses, PPARγ does not appear to mediate the anti-inflammatory effects of nitro-fatty acids in macrophages.^7^ Nitro-fatty acids were shown to inhibit type I interferon (IFN) release by macrophages, independently of PPARγ and KEAP1-NRF2 pathways, through direct nitro-alkylation of Stimulator of interferon genes (STING).^8^ What other cellular targets or signaling pathways are targeted by nitro-fatty acids? Are there molecular and structural determinants for protein nitro-alkylation? Characterization of proteomes that are susceptible to nitro-alkylation is critical for elucidating modes of action and cellular functions of nitro-fatty acids.

Here we aim to characterize the protein targets of nitro-fatty acids in living cells using a chemoproteomics strategy. We synthesized, characterized and applied a clickable nitro-oleate probe for the detection and global identification of nitro-alkylated proteins in mammalian cells. 184 nitro-alkylated proteins were identified in human THP1 macrophages, including known nitro-alkylated proteins like KEAP1 and STING. Transmembrane proteins, especially those localized to endoplasmic reticulum (ER) and nuclear membranes, were enriched in nitro-alkylated proteins. Nitro-alkylated proteins were also significantly enriched in lipid metabolism and transport proteins. This study provides the first global profile of protein targets of nitro-fatty acids in living cells and serves as a useful resource to understand the pleiotropic biological activities of nitro-fatty acids.

## 2. Materials and methods

### 2.1. Chemicals and probes

All chemicals and reagents were purchased from Sigma-Aldrich unless otherwise indicated. Azide-rhodamine was synthesized as previously described.^9^ Diazo-biotin-azide was purchased from DC Biosciences. Alkyne-oleic acid, oleic acid and 9-NO_2_-OA were purchased from Cayman Chemicals. For detailed alk-9-NO_2_-OA synthesis and characterization, refer to Supporting information.

### 2.2. Cell lines

Hek293T and Hek293FT cells were maintained in Dulbecco’s High Glucose Modified Eagle’s Medium (DMEM, Hyclone) containing 10% heat-inactivated fetal bovine serum (FBS, Biological Industries), 100 U/mL penicillin and 100 μg/mL streptomycin (Gibco). HEK-Blue IFN-α/β cells (InvivoGen) were propagated in DMEM with 10% FBS, 100 μg/mL normocin, 30 μg/mL blasticidin and 100 μg/mL zeocin (InvivoGen). THP1, Jurkat, Raji, and Ramos cells were cultured in Roswell Park Memorial Institute (RPMI, Hyclone) 1640 supplemented with 10% FBS. THP1 monocytes were differentiated into adherent macrophages by 48 h incubation with 200 nM phorbol 12-myristate 13-acetate (PMA) followed by 48 h rest in PMA-free medium. All cells were maintained at 37 °C in a humidified incubator with 5% CO_2_.

### 2.3. Type I IFN assays

PMA-differentiated THP1 macrophages in 24-well plates were pretreated with fatty acids (2.5 μM and 10 μM) for 30 min prior to stimulation with 4 μg/mL cGAMP (InvivoGen) using Lipofectamine 2000 (Invitrogen). After 20 h stimulation, culture supernatants were analyzed for human type I IFN activity using the HEK-Blue IFN-α/β reporter cell line according to the manufacturer’s instructions.

### 2.4. Cell viability assay

Cell viability was measured using the colorimetric MTS assay (Promega) according to manufacturer’s instructions. Briefly, after removing culture supernatants, cells were washed with PBS and 250 μL of the MTS-PMS mixture solution was added. The cells were incubated for 0.5-1.5 h at 37 °C before absorbance at 490 nm was measured in a microplate reader. Data was normalized using cells untreated with compounds (100%) and buffer only background controls (0%).

### 2.5. Wild type and mutant KEAP1 overexpression constructs

The following mutations to pCMV3-HA-KEAP1 (Sino Biological) were made using the QuikChange Lightning Multi Site-Directed Mutagenesis Kit (Stratagene) and validated by Sanger sequencing. 3M: C151S, C273W, C288E. 7M: C38S, C151S, C226S, C257S, C273W, C288E, C489S. These constructs were transfected into HEK293T cells using the TransIT-LT1 reagent (Mirus Bio) for 24 h prior to treatment with fatty acids and alk-9-NO_2_-OA.

### 2.6. Fluorescent detection of nitro-fatty acid protein targets in living cells

Cells were grown to ~90% confluence prior to treatment with clickable chemical probes (alk-OA or alk-9-NO_2_-OA) at the indicated concentrations with the same volumes of DMSO or indicated fatty acids used as controls. At the described time points, labelled cells were washed thrice with ice-cold PBS and pelleted at 400 *g* for 5 min. Cells were flash-frozen in liquid nitrogen and stored at −80 °C prior to lysis. Frozen cells were lysed in SDS lysis buffer [4% SDS, 150 mM NaCl, 50 mM triethanolamine, pH 7.4, 2× EDTA-free protease inhibitor cocktail (Thermo Scientific), 10 mM phenylmethylsulfonyl fluoride, 50 U/mL SuperNuclease (Sino Biological)]. Protein concentrations were determined by BCA protein assay (Thermo Scientific). For in-gel fluorescence detection of nitro-fatty acid protein targets, 50 μg cell lysates were reacted with freshly made CuAAC reaction cocktail [100 μM azide-rhodamine, 1 mM CuSO_4_, 1 mM tris(2-carboxyethyl) phosphine hydrochloride (TCEP), 100 μM tris[(1-benzyl-1H-1,2,3-triazol-4-yl)methyl]amine (TBTA)] in a total reaction volume of 50 μL for 1 h at room temperature. Proteins were chloroform-methanol precipitated and the protein pellet was washed twice with ice-cold methanol. Air-dried protein pellets were resuspended in 25 μL SDS buffer before the addition of 8.7 μL 4× SDS-loading buffer (20% glycerol, 125 mM Tris·HCl, pH 6.8, 4% SDS, 0.05% bromophenol blue) and 1.3 μL Bond-Breaker TCEP (Thermo Scientific). Samples were heated for 5 min at 75 °C, separated by SDS-PAGE, and imaged on a Typhoon 9400 variable-mode imager. Rhodamine-associated signal was detected at excitation 532 nm/emission 580 nm. After fluorescence scanning, gels were either stained with Coomassie (Protein Ark) or transferred to PVDF membranes for Western blot analysis.

### 2.7. Antibodies used for Western blot analysis

The following primary and secondary antibodies were used for Western blot analysis in this study. Anti-HA (#05-904) and anti-ESYT2 (HPA002132) were purchased from Sigma. Anti-GAPDH (#5174), anti-STING (#13647), anti-NR3C1 (#12041), anti-STAT3 (#12640), anti-TLR2 (#12276) and anti-RXRα (#3085) were purchased from Cell Signaling Technology. Streptavidin-HRP (#7043) was from Abcam. Anti-mouse-HRP (#115-035-003) and anti-rabbit-HRP (#111-035-003) were purchased from Jackson ImmunoResearch Laboratories.

### 2.8. Affinity enrichment of nitro-alkylated proteins

For affinity purification of alk-9-NO_2_-OA modified proteins, 400 μg of cell lysates were diluted with HEPES buffer (150 mM NaCl, 50 mM HEPES pH 7.4) to 360 μL, and 40 μL of freshly prepared CuAAC reaction cocktail [diazo-biotin-azide (4 μL, 10mM stock solution in DMSO), CuSO_4_ (8 μL, 50 mM aqueous solution), TCEP (8 μL, 50 mM aqueous solution), TBTA (20 μL, 2 mM stock solution in DMSO)] was added. After 1 h incubation at room temperature, EDTA (8 μL of 0.5 M solution) was added to stop the CuAAC reaction and samples were chloroform-methanol precipitated. Protein pellets were allowed to air-dry, resuspended in 80 μL SDS-HEPES buffer (4% SDS, 150 mM NaCl, 50 mM HEPES pH 7.4), diluted to 0.5 mg/mL with HEPES buffer. The protein samples were then added to 20 μL of high-capacity NeutrAvidin beads (Thermo Scientific), and incubated with end-over-end rotation for 90 min at room temperature. The beads were sequentially washed with 1 mL of 1% SDS in PBS (3×5 min), 4 M urea in PBS (2×5 min) and AMBIC (50 mM ammonium bicarbonate) (5×2 min). For SDS-PAGE and western blot analysis, 25 μL of freshly prepared elution buffer (25 mM Na_2_S_2_O_4_ in PBS with 0.1% SDS) were added to the beads and the samples were eluted by incubation for 1 h at room temperature. For LC-ESI MS/MS analysis, samples were reduced with 10 mM TCEP (stock solution 100 mM pH 8 in AMBIC) for 30 min at room temperature. After removing the supernatant and washing the beads once with AMBIC, samples were incubated with 10 mM iodoacetamide (stock solution 100 mM pH 8 in AMBIC) at room temperature in the dark for 30 min. The supernatant was removed and the beads were washed twice with AMBIC. Finally, 100 μL of AMBIC containing trypsin (0.1 μg trypsin/500 μg lysate) was added for on-bead digestion overnight at 37 °C. Samples were centrifuged and the supernatant was transferred into clean tubes. The beads were washed with 100 μL of 1% formic acid (FA)/15% acetonitrile (ACN) in H_2_O, followed by 100 μL of 1% FA in H_2_O. These washes were combined with the supernatant, and the peptides were concentrated and desalted using C18 spin tips (Thermo Scientific) according to the manufacturer’s instructions. Peptides were dried in the speedvac and resuspended prior to LC-ESI MS/MS analysis.

### 2.9. LC-ESI MS/MS analysis by Orbitrap-MS

Peptides extracted from the on-bead digested samples were analyzed by the LTQ-Orbitrap mass spectrometer (Thermo Fisher) coupled with nano-LC (Dionex Ultimate 3000; Thermo Fisher) for protein identification. Dried peptides were reconstituted in buffer A (0.1% FA) and injected into the trap column (Zorbax 300SB-C18, 0.3 5 mm; Agilent Technologies) at a flow rate of 0.18 μL/min in buffer A. The peptide mixture was fractionated using a nano-C18 resolving column (inner diameter 75 μm; column length 10 cm) for electrospray ionization using a 15 μm tip (New Objective). A 70 min gradient time was used to separate the peptides for LC-MS/MS analysis. The peptides were eluted using a linear gradient of 0–8% HPLC buffer B (99.9% ACN containing 0.1% FA) for 4.5 min, 8–16% buffer B for 0.5 min, 16–45% buffer B for 38 min, 45-60% buffer B for 10 min, 60–80% buffer B for 5 min, 95% buffer B for 5 min, and 8% buffer B for 7 min at a flow rate of 0.25 μL/min. Xcalibur 2.0 software (Thermo Fisher) was used for the LTQ-Orbitrap operation. Twenty data-dependent MS/MS scan events (in the linear ion trap) were followed by one MS scan for the ten most abundant precursor ions in the preview MS scan at a resolution of 120,000. Full-scan MS was performed in the Orbitrap over a range of 400 to 1,450 Da. The (Si(CH3)2O)6H+ ion signal at m/z 445.120025 was used for internal calibration of accurate mass accuracy.

### 2.10. MS data processing, database search for protein identification

Mass spectra were processed and searched using Proteome Discoverer (Version 1.4.1.14, Thermo Fisher) against Swiss-Prot human database (released Jan, 2020, 20368 entries of *Homo sapiens)* of the European Bioinformatics Institute. The mass tolerances for parent and fragment ions were set as 10 ppm and 0.5 Da. The carbamidomethylation on cysteine (+57.0 Da) was set as fixed modification. Oxidation on methionine (+15.99 Da) was set as variable modifications. The enzyme was set as trypsin and up to two missed cleavage was allowed. The protein identification threshold was set as 0.95 to ensure an overall false discovery rate below 0.5%. Proteins with at least one unique peptide identified were retained in this study.

### 2.11. Bioinformatic analysis of high-confidence nitro-alkylated protein hits

GO enrichment analysis was performed using the DAVID.^10,11^ After Benjamini-Hochberg correction for multiple testing, Cellular Component GO terms with adjusted p-values (q-values) <0.05 were selected. Information on topology of membrane proteins (multi-pass, single-pass, non-transmembrane) of nitro-alkylated protein hits and the *H. sapiens* reference genome was extracted from Swiss-Prot (accessed 2021-05-20). Pathway and process enrichment analysis was performed using Metascape with the following ontology sources: KEGG Pathway, GO Biological Processes, Reactome Gene Sets, Canonical Pathways, CORUM, TRRUST, DisGeNET, PaGenBase, Transcription Factor Targets, WikiPathways and COVID.^12^ Only the top 20 functional clusters with Benjamini-Hochberg corrected p-values (q-values) <0.05 were selected. A hierarchically clustered binary heatmap showing the common protein targets of nitro-fatty acid and other electrophiles (Supplementary Table 2) was generated using Morpheus.^13^

### 2.12. Glucocorticoid receptor radioligand binding assay

Assays were performed at Eurofins Scientific. Briefly, full-length human recombinant glucocorticoid receptor expressed in insect cells was used in modified potassium phosphate buffer pH 7.4. For each data point, a 10 μg aliquot was incubated with 5 nM [^3^H]dexamethasone, 1 μg anti-GST antibody and 0.2 mg scintillation proximity assay (SPA) beads for 24 h at 4 °C. Non-specific binding was estimated in the presence of 10 μM dexamethasone. Receptors were counted to determine specifically bound [^3^H]dexamethasone. Compounds were screened at 10 μM. Assays were conducted in duplicates and ≥50% of maximum inhibition was considered significant.

### 2.13. Docking analysis of 9-NO_2_-OA with glucocorticoid receptor-ligand binding domain

The protein structure of the NR3C1 ligand-binding domain (NR3C1-LBD) (Protein Data Bank identifier (PDB ID: 1NHZ) was used for this study.^14^ We performed the two-step method to get the covalent binding of ligand and NR3C1-LBD. 9-NO_2_-OA was first docked into the binding site by BIOVIA 2018/LigandFit program (BIOVIA, Inc., San Diego, CA).^15^ The binding pocket was identified from the co-crystal structure binding site of antagonist RU-486 (PDB ID: 1NHZ). The piecewise linear potential 1 (PLP1) force field for calculating ligand-receptor interaction energies was used. The default grid parameters were set with 0.5 Å spacing and the extension from Site was set as 7 Å. The number of docking poses was set as 100 conformations and the docking RMS threshold for ligand-site matching was set as 5 Å. The steepest descent steps minimization with CHARMM force field was performed as the compound dock into the binding site.^16^ Finally, the covalent docking calculation was performed to illustrate the binding of 9-NO_2_-OA. The docking was conducted by using two-point attractor method to form the covalent bonding by the AutoDock Tools (version 1.5.6).^17^ The number of docking poses was set to 20 with default parameters. The decision of the best pose was according to the lowest binding energy of the 9-NO_2_-OA, which forms a covalent bond with Cys643 of NR3C1-LBD.

## 3. Results

### 3.1. Regio- and stereo-selective synthesis of a clickable nitro-oleic acid probe

Chemoproteomic approaches with chemical probes have emerged as invaluable tools to profile the cellular targets of bioactive small molecules, including metabolites, natural products and drugs in a variety of biological systems.^18^ In particular, clickable lipid probes harboring the relatively small terminal alkyne group retained their ability to modify proteins while allowing subsequent installation of fluorescence or affinity tags through copper-catalyzed azide-alkyne cycloaddition (CuAAC) for the detection and enrichment of lipid-modified proteins.^19^ Therefore, to detect and profile nitro-alkylated proteins in native biological systems, we synthesized a clickable nitro-fatty acid probe. We chose to install the alkyne group at the terminal carbon away from the carboxylic acid group, which is critical for the formation of coenzyme A thioesters during fatty acid trafficking and metabolism. Since positional and geometric isomers of nitro-oleate can have different substrate reactivities or biological activities,^20,21^ we sought to regio- and stereo-selectively synthesize the alkynyl-(*E*)-9-nitro-oleic acid (alk-9-NO_2_-OA) probe. Among the different approaches to synthesize natural nitro-fatty acids and their derivatives,^22–25^ we selected one employing the nitro-aldol condensation reaction, in which the regiochemistry of the nitro group is fixed by the precursor combination (Scheme 1).^22^ 9-Bromononanol was first oxidized to carboxylic acid **1** and after protection, the bromo moiety of *tert-butyl* ester **2** was transformed to a nitro moiety to produce **3**. Following a two-step conversion of undec-10-ynoic acid to aldehyde **5**, nitro-aldol condensation of **3** and **5** yielded a β-hydroxynitro ester **6** as 45:55 mixture of diastereomers. Finally, the desired probe **8** was synthesized in three steps via acetylation of the hydroxyl group, base-induced elimination and deprotection. The stereochemistry of **8** was confirmed by ^1^H-NMR comparison with the reported (*E*)-nitroalkene.^22^ This synthetic strategy can potentially be used to regio- and stereo-selectively synthesize other clickable nitro-fatty acid probes to complement structure-activity relationship studies.

### 3.2. Fluorescent detection of nitro-alkylated proteins

To establish alk-9-NO_2_-OA as a chemical reporter for protein nitro-alkylation, we evaluated its ability to enter living cells and modify proteins, including known nitro-alkylated proteins. HEK293FT cells were treated with varying concentrations of alk-9-NO_2_-OA for different lengths of time. To determine if alk-9-NO_2_-OA will modify a known nitro-alkylated protein, HEK293FT cells overexpressing HA-KEAP1 were similarly treated. The cell lysates were reacted with azide-rhodamine by CuAAC prior to separation by SDS-PAGE (Figure 1A). Ingel fluorescence revealed that alk-9-NO_2_-OA could enter living cells and label intracellular proteins such as KEAP1. Rapid labelling of proteins by alk-9-NO_2_-OA was observed within minutes (Figure 1B), which is unsurprising given the significantly higher thiol reaction rates observed for nitro-fatty acids compared to non-nitrated electrophilic lipids.^5^ While there was dose- and time-dependent labeling of KEAP1 and other endogenous proteins by alk-9-NO_2_-OA, probe concentration had a greater effect on labelling intensity (Figure 1C). Protein labeling by alk-9-NO_2_-OA was specific, as demonstrated by decreased labelling of KEAP1 when multiple known OA-NO_2_-reactive cysteines were mutated (Figure 1D).^26^ Gel-based profiling with alk-9-NO_2_-OA in different human cell lines revealed both common and distinct cellular targets that potentially contribute to the wide range of biological activities observed for nitrofatty acids (Figure 1E).

**Figure 1.**
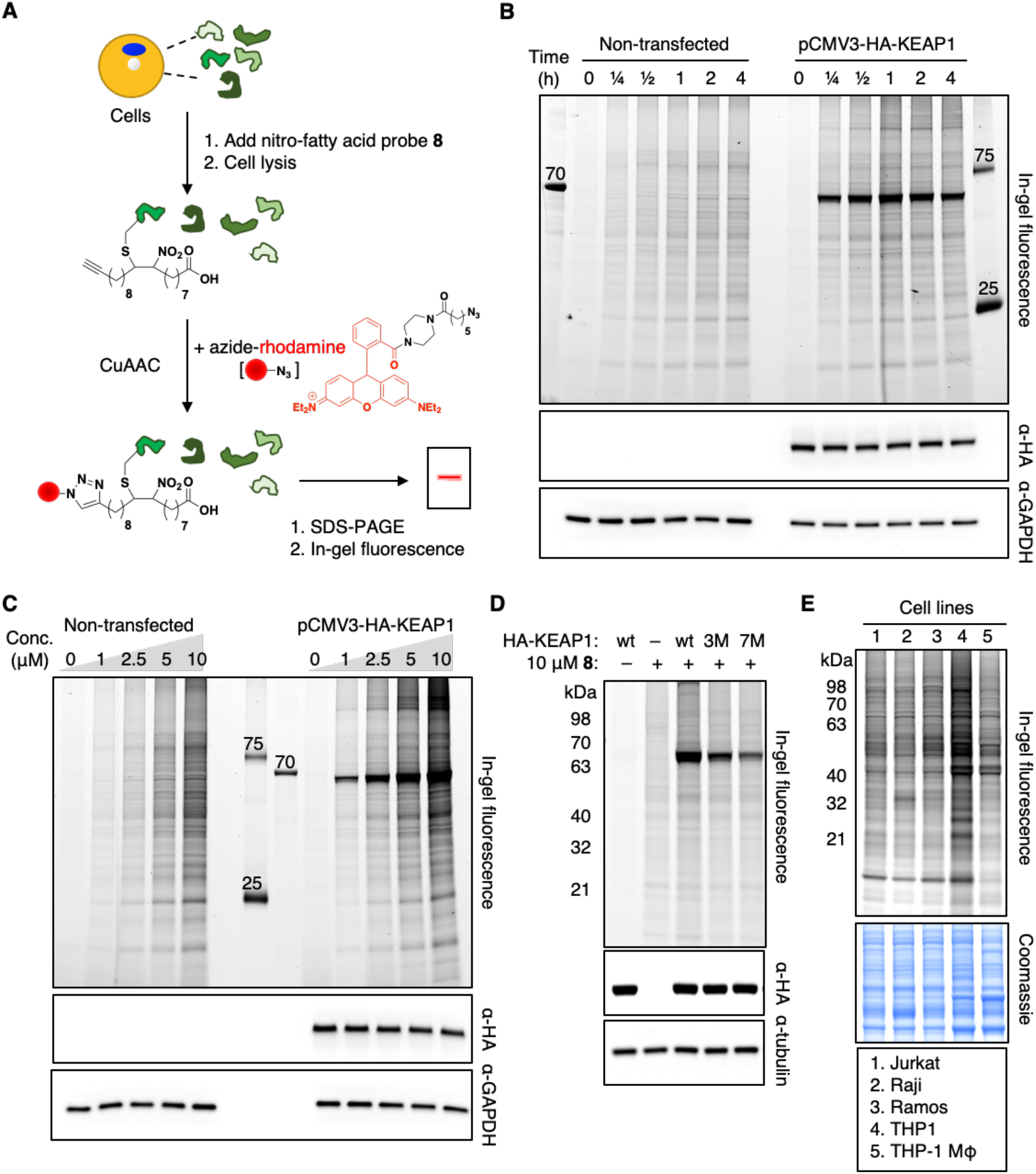
Fluorescent detection of nitro-alkylated proteins in living cells. **(A)** Workflow for in-gel fluorescence detection of nitro-alkylated proteins using the clickable alk-9-NO_2_-OA probe **8** and azide-rhodamine. CuAAC, copper-catalyzed azide-alkyne cycloaddition. **(B)** Time-dependent labelling of endogenous proteins and overexpressed HA-KEAP1 in HEK293FT cells at 2.5 μM alk-9-NO_2_-OA. **(C)** Dose-dependent labelling of endogenous proteins and overexpressed HA-KEAP1 in HEK293FT cells by alk-9-NO_2_-OA. **(D)** Labeling of wildtype (wt) KEAP1 and its mutants in HEK293FT cells at 2.5 μM alk-9-NO_2_-OA. 3M and 7M refer to mutants with 3 and 7 cysteines mutated, respectively. **(E)** Fluorescent detection of nitro-alkylated proteins in different human cell lines. Anti-HA, anti-GAPDH, anti-tubulin blots and Coomassie stains act as loading controls for the accompanying fluorescence gels. Selected protein molecular weight markers are indicated for each gel.

### 3.3. Selective enrichment of nitro-alkylated proteins in THP1 macrophages

To globally profile nitro-alkylated proteins as well as uncover novel cellular targets of nitro-fatty acids, we chose to work in THP1 macrophages. This system, in which nitro-fatty acids have been shown to inhibit type I IFN response by alkylating and inhibiting STING,^8^ provides an excellent model biological system to functionally validate our probe and explore the scope of protein nitro-alkylation in living cells.

We first confirmed that the alk-9-NO_2_-OA probe retained the ability to inhibit STING function in cells despite harboring an additional alkyne group compared to the original nitro-fatty acid. Consistent with the study by Hansen and colleagues,^8^ 9-NO_2_-OA and alk-9-NO_2_-OA significantly inhibited type I IFN response upon STING stimulation with cyclic guanosine monophosphate-adenosine monophosphate (cGAMP) without affecting cellular viability at 10 μM (Figure 2A). This inhibitory effect was dependent on the nitro group of alk-9-NO_2_-OA since type I IFN release was unaffected by alk-OA.

**Figure 2.**
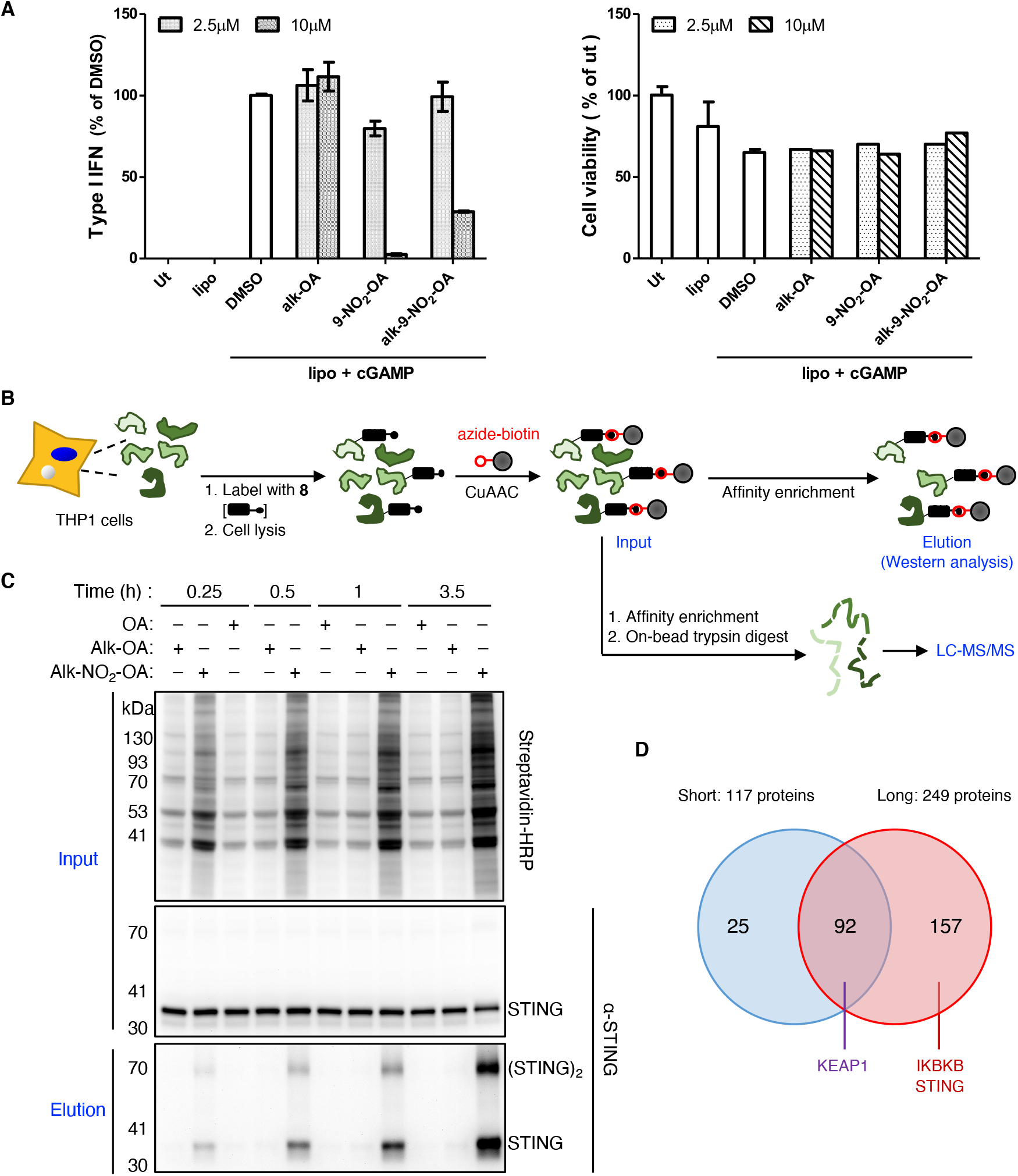
Selective enrichment and identification of nitro-alkylated proteins in living THP1 macrophages. **(A)** PMA-differentiated THP1 macrophages were treated with the indicated fatty acids (2.5 μM and 10 μM) for 30 min before stimulation with cGAMP using Lipofectamine 2000 (Lipo). After 20 h, culture supernatants analyzed for type I IFN production (left panel) relative to the DMSO control (left panel) and cell viability (right panel) relative to the untreated control (ut). Error bars, standard deviation. **(B)** Enrichment scheme for nitro-alkylated proteins using the clickable alk-9-NO_2_-OA probe **8** and diazo-biotin-azide (azide-biotin). CuAAC, copper-catalyzed azide-alkyne cycloaddition. Enriched proteins can be analyzed by western blot or LC-MS/MS analysis. **(C)** Prior to affinity purification, protein samples (input, top panels) labeled with the indicated lipids for different times were analyzed using streptavidin and anti-STING antibodies. STING was selectively recovered from THP1 cells labeled with the alk-9-NO_2_-OA (elution, bottom panel). **(D)** Number of candidate nitro-alkylated proteins identified from cells treated with 10 μM alk-9-NO_2_-OA for 15 min (short) and 3.5 h (long) from three biological repeats as well as overlap between the two treatment conditions. Known nitro-alkylated proteins are indicated. For proteomics filter criteria, see Supplementary Figure 2 and Methods.

We further showed that alk-9-NO_2_-OA formed a protein adduct with endogenous STING in THP1 macrophages by reacting cell lysates with azide-functionalized biotin and demonstrating selective enrichment of STING using NeutrAvidin (Figure 2B and 2C). STING alkylation and enrichment required the nitro moiety of alk-9-NO_2_-OA since no enrichment of STING was observed in samples treated with alk-OA. Similar STING levels in the input samples ruled out differences in protein expression. Interestingly, for alk-9-NO_2_-OA treated samples, we observed an enrichment of dimeric STING species (Figure 2C), which has been reported to resist complete dissociation and exhibit slower mobility in SDS-PAGE.^27,28^ This suggested that nitro-alkylated STING may be potentially more capable of retaining its dimeric conformation or that dimeric STING may be preferentially nitro-alkylated.

Collectively, these experiments indicated that alk-9-NO_2_-OA probe retained its biological activity to suppress type I IFN response in macrophages and can be used to selectively enrich a known nitro-alkylated protein expressed at endogenous levels.

### 3.4. Global profiling of nitro-alkylated proteins in THP1 macrophages

Given that the alk-9-NO_2_-OA probe can be used to selectively label and enrich known nitro-alkylated proteins, we proceeded to explore the scope of protein nitro-alkylation by profiling nitro-alkylated proteins in living cells using a chemoproteomics strategy (Figure 2C). THP1 macrophages were treated with 10 μM alk-9-NO_2_-OA for 15 min or 3.5 h. As specificity controls, macrophages were similarly treated with 9-NO_2_-OA and alk-OA. After cell lysis, modified proteins were conjugated to biotin by CuAAC, enriched using NeutrAvidin, digested on-bead with trypsin, and analyzed by LC-MS/MS.

117 and 249 proteins identified in at least two of three “15 min” or “3.5 h” biological repeats and were absent in the 9-NO_2_-OA samples constituted our list of “short” and “long” labeled nitro-alkylated proteins respectively (Figure 2D). The identification of established nitro-alkylated proteins, KEAP1, STING and Inhibitor of nuclear factor kappa-B kinase subunit beta (IKBKB),^8,26,29^ further demonstrated that our probe and strategy is capable of capturing physiological targets of nitro-fatty acids in living cells. The identification of STING only in “long” samples was consistent with our previous western blot analysis that showed increasing STING enrichment with longer treatment times (Figure 2C). Notably, we did not identify any fatty acid binding proteins as our approach identifies proteins that form covalent adducts with nitro-fatty acids. Since most of the “short” label list of nitro-alkylated protein list were also found in the “long” label list (Figure 2D), we chose to not to differentiate between the two treatment conditions for subsequent bioinformatic analyses. After selecting proteins that were absent in 9-NO_2_-OA samples and found in at least three of the six alk-9-NO_2_-OA samples (Supplementary Figure 2), subsequent exclusion of proteins that were found in alk-OA samples yielded a high-confidence list of 184 nitro-alkylated protein hits (Supplementary Table 1), majority of which are not known to be targets of nitro-fatty acids.

### 3.5. Bioinformatic analysis of nitro-alkylated protein hits

Enrichment analyses were performed to search for features or functions that are enriched in nitro-alkylated proteins. Gene ontology (GO) analysis of our nitro-alkylated protein hits revealed that exogenously added nitro-fatty acids were able to enter cells and modify intracellular proteins. ER and nuclear membrane proteins as well as mitochondrial and peroxisomal proteins were significantly enriched with nitro-alkylated proteins (Figure 3A), suggesting that nitro-fatty acids may be enriched in endomembrane compartments involved in lipid metabolism. This was further reflected by the enrichment of transmembrane proteins in our nitro-alkylated protein list compared to the human reference proteome (Figure 3B). Pathway and process analysis revealed that nitro-alkylated proteins were overrepresented in lipid metabolism, lipid transport, maintenance of location, one-carbon metabolism, nuclear receptor meta-pathways, among others (Figure 3C).

**Figure 3.**
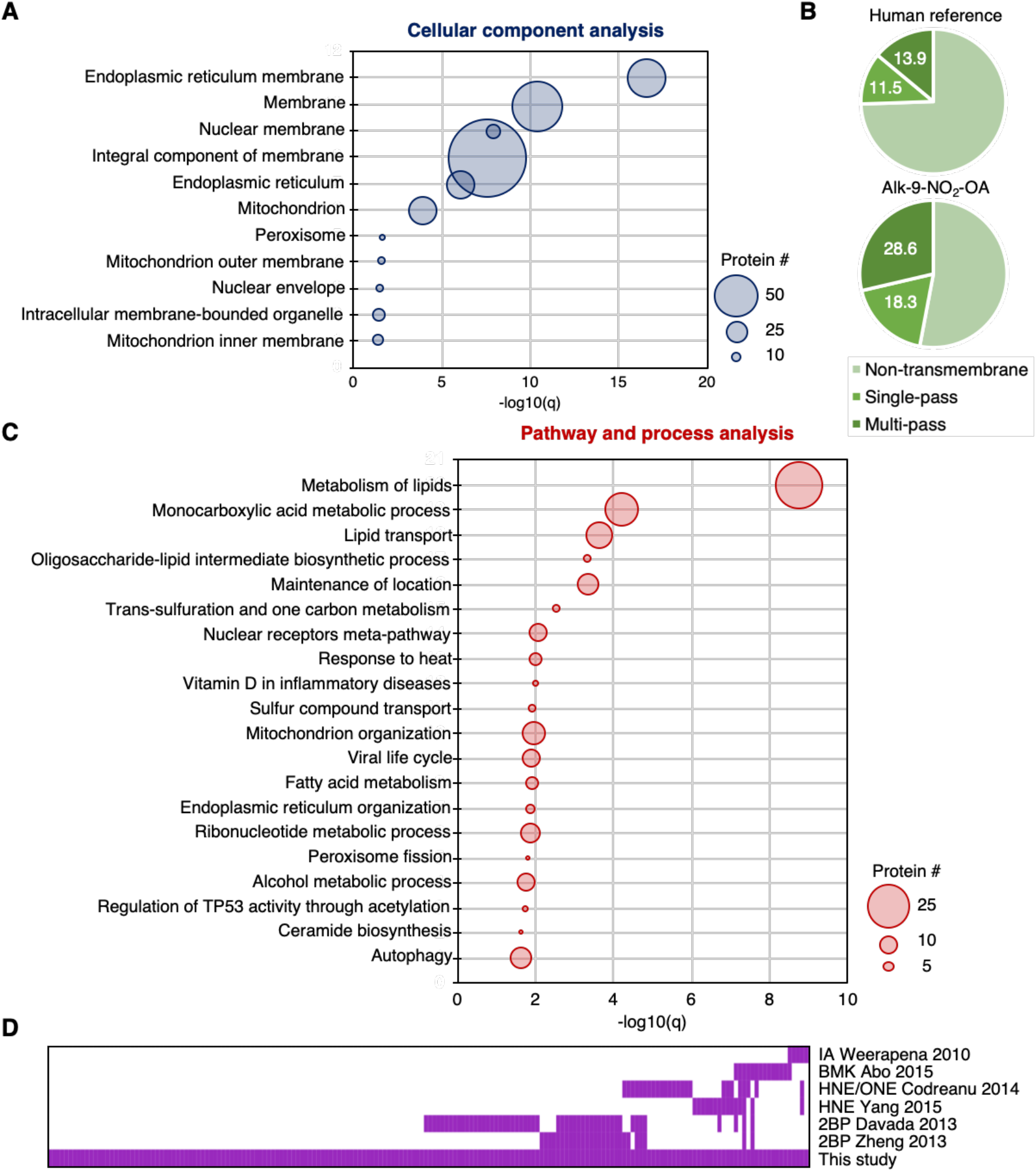
Bioinformatics analysis of 184 high-confidence nitro-alkylated proteins. **(A)** GO cellular component analysis of candidate nitro-alkylated proteins. Cellular components and their Benjamini-adjusted p-values (q-value) of more than 1.3 are shown. Size of each circle reflects the number of the protein associated with each cellular component. **(B)** Topology analysis of candidate nitro-alkylated protein and the human reference proteome based on UniProt annotation. **(C)** Metascape pathway and process analysis of candidate nitro-alkylated proteins. The top 20 clusters and their q-values are shown and size of each circle reflects the number of the protein associated with each cluster. **(D)** Comparative analysis of the protein targets of nitro-fatty acids and other electrophiles. Each column represents one of the 184 high confidence nitro-alkylated proteins and proteins found to be modified by the indicated electrophiles in their respective studies are marked purple (Supplementary Table 2). IA, iodoacetamide.^32^ BMK, bromomethyl ketone.^33^ HNE, 4-hydroxy-2-nonenal and ONE, 4-oxo-2-nonenal.^47,48^ 2BP, 2-bromopalmitate.^30,31^

To gain insight into the reactivity and substrate selectivity of nitro-fatty acids, we performed a comparative analysis of nitro-alkylated proteins with proteins modified by other electrophiles, which include naturally occurring lipid-derived aldehyde electrophiles HNE and 4-oxo-2-nonenal, as well as a non-selective inhibitor of lipid metabolism 2-bromopalmitate.^30,31^ In addition, since thiol-reactive iodoacetamide and bromomethylketone probes have been used to broadly profile and monitor reactive functional cysteine residues that are involved in redox regulation, post translational modification, catalysis and protein recognition,^32,33^ we examined if our list of nitro-alkylated proteins contained these “reactive” cysteines. Interestingly, while there was limited overlap, about half of our high-confidence nitro-alkylated proteins are not targeted by the other electrophiles that we looked at (Figure 3D). The latter group of proteins, which includes STING and IKBKB (Supplementary Table 2), may contribute to the unique regulatory roles observed for nitro-fatty acids in various inflammatory and fibrotic disease models.

### 3.6. Validation of novel nitro-alkylated proteins

To confirm our LC-MS/MS results, we selected five proteins identified with a range of spectral counts and different cellular localization for further validation by western blot (Figure 4A). As expected for nitro-alkylated proteins, all five proteins, Extended synaptotagmin 2 (ESYT2), Signal transducer and activator of transcription 3 (STAT3), Toll-like receptor 2 (TLR2), Retinoid X receptor alpha (RXRα) and Glucocorticoid receptor (NR3C1), were selectively enriched in samples exposed to alk-9-NO_2_-OA but not in those treated with control lipids, 9-NO_2_-OA and alk-OA (Figure 4B). Similar levels of each target protein prior to enrichment ruled out discrepancies in protein expression due to the different lipid treatments.

**Figure 4.**
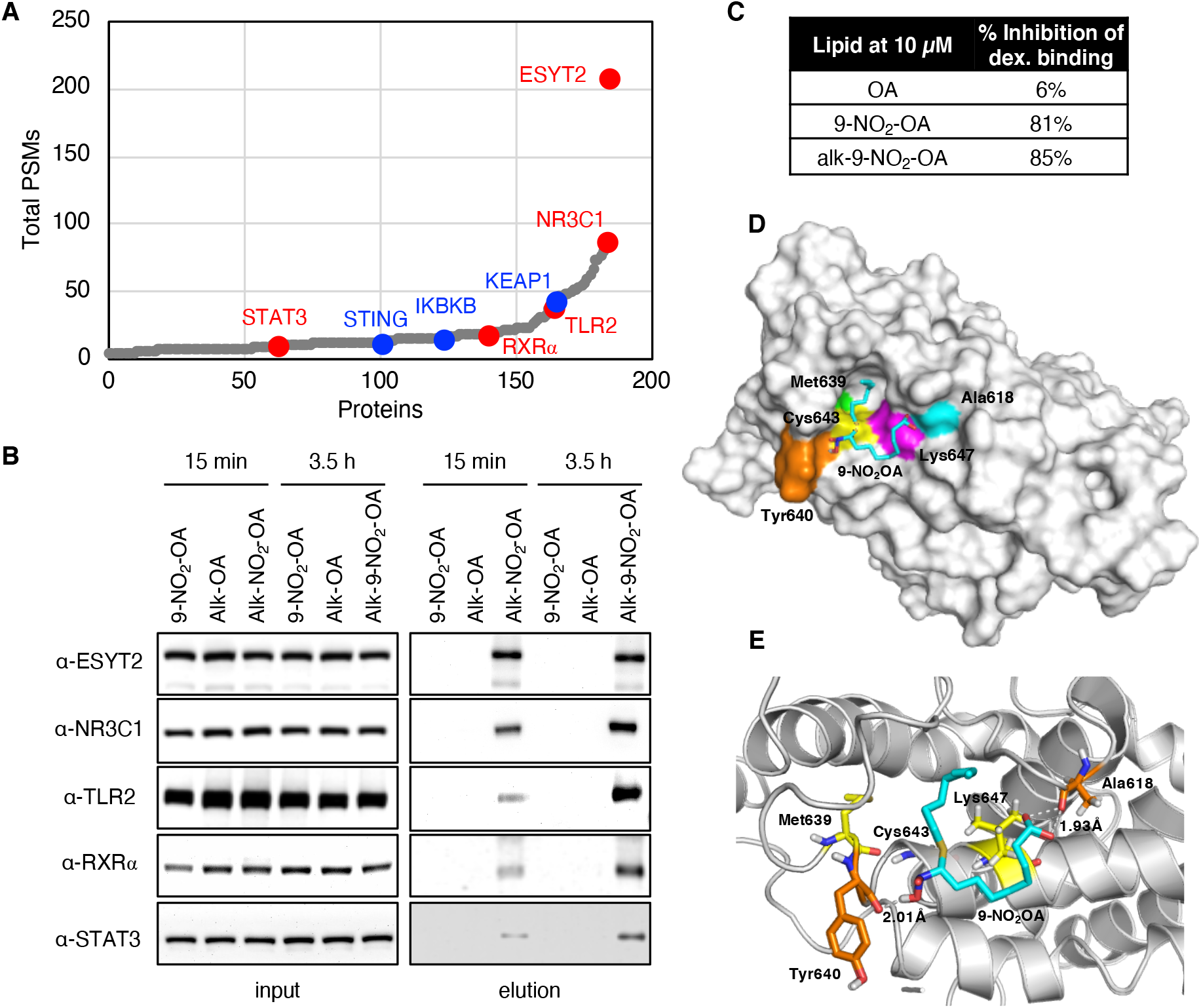
Validation of novel nitro-alkylated proteins. **(A)** 184 proteins in our high-confidence list nitro-alkylated proteins were sorted with increasing total number of PSMs detected for alk-9-NO_2_-OA samples in the chemoproteomics study. Characterized and novel nitro-alkylated proteins validated in this study are highlighted in red and blue, respectively. Nitro-alkylation of STAT3 was demonstrated in a separate study during the preparation of this manuscript.^39^ **(B)** Lysates from cells were treated with the indicated lipids for 15 min or 3.5 h were reacted with diazo-biotin-azide via CuAAC. Protein samples before NeutrAvidin enrichment (input) and after elution were immunoblotted for the indicated target proteins. **(C)** NR3C1 radioligand binding assay. The percent inhibition of [^3^H]dexamethasone binding to full-length human NR3C1 in the presence of 10 μM lipid is shown. **(D)** The most stable complex obtained for 9-NO_2_-OA in the NR3C1-LBD (1NHZ). 9-NO_2_-OA and relevant residues are colored. **(E)** Binding site analysis for 9-NO_2_-OA (Cyan) with NR3C1-LBD. 9-NO_2_-OA forms a covalent bond with Cys643, and hydrogen bonds with Tyr64O and Ala618 (orange residues). Hydrophobic effect was contributed by Leu647 and Met639 (yellow residues).

To further examine the effect of nitro-alkylation on NR3C1 function, we tested the ability of different lipids to inhibit radiolabeled dexamethasone binding to full-length human NR3C1 *in vitro*. Both 9-NO_2_-OA and alk-9-NO_2_-OA significantly inhibited dexamethasone binding to NR3C1 at 10 μM compared to OA, which had no effect on radioligand binding (Figure 4C). We further performed two-step molecular docking was performed to investigate the covalent binding interactions of 9-NO_2_-OA within the NR3C1-LBD (Figure 4D). 9-NO_2_-OA is predicted to form a covalent bond with Cys643 and stabilized within the glucocorticoid binding pocket by a strong binding network by hydrogen bonding with Tyr640 and Ala618 and hydrophobic interactions with Leu647, and Met639 (Figure 4E), with a calculated interaction energy of −136.51 kcal/mol. These results suggested that 9-NO_2_-OA potentially affects NR3C1 ligand binding ability via nitro-alkylation.

## 4. Discussion

Our global characterization of the mammalian proteome susceptible to alkylation by 9-NO_2_-OA in living cells identifies 184 proteins, majority of which are not known to be targets of nitro-fatty acids. The overrepresentation of nitro-alkylated proteins in endomembrane compartments involved in lipid metabolism, in particular ER/nuclear membrane and mitochondria, suggest possible enrichment of the nitro-fatty acid these compartments. This is perhaps unsurprising given that nitro-fatty acids are actively metabolized *in vivo* via β-oxidation or esterified and incorporated into glycerophospholipids and triacylglycerols.^3,21,34,35^ Nonetheless, nitro-alkylation of lipid metabolism and transport proteins in these membrane compartments suggests that nitro-fatty acids directly modulate cellular lipid metabolism (*vide infra*). Comparative analysis between proteins that are susceptible to modification by different endogenous and synthetic electrophiles offer insight into the unique biological activities observed for nitro-fatty acids. Given the noticeably different profiles of nitro-alkylated proteins in different cell lines, our list is most likely an underestimate of the full complement of proteins that are susceptible to context-dependent alkylation and modulation by nitro-fatty acids.

Many studies on the biological effects of nitro-fatty acids focused on a few well-characterized protein targets and their associated pathways. While established nitro-alkylated proteins like STING, KEAP1 and IKBKB were identified in our study, others like PPARγ were noticeably absent. This may be due to the differences in the chemical structures of the nitro-fatty acid, cellular states or detection methods. Compared to regioisomer 10-NO_2_-OA, 9-NO_2_-OA is more reactive towards KEAP1 cysteines,^26,36^ but is less reactive in the nitro-alkylation of PPARγ.^37^ Bervejillo et al. showed that nitro-fatty acids are less able to activate PPARγ in macrophages compared to monocytes,^7^ suggesting that PPARγ may not be a major target for nitro-fatty acids in macrophages. Our discovery that important signaling molecules like STAT3, TLR2, RXRα and NR3C1 can be nitro-alkylated indicates a much broader regulatory role of nitro-fatty acids in inflammation and immune modulation than previously appreciated. Indeed, broad effects of nitro-oleate on both pro- and anti-inflammatory macrophage functions have been observed.^38^ Notably, while we were preparing this manuscript, Wang et al. reported that nitro-alkylation of STAT3 inhibits its phosphorylation and activation, thereby contributing to the anti-inflammatory role of nitro-oleate.^39^ This study supports that our chemical probe and chemoproteomic approach can be used to uncover novel nitro-alkylated proteins. Overall, our study demonstrates the multi-target pharmacology of nitro-fatty acids where in macrophages, key regulatory proteins in TLR, JAK-STAT, NF-κB, nuclear hormone receptor and redox signaling pathways that are central to macrophage function are targeted.

Nitro-fatty acids play important roles in modulating lipid metabolism in cells and animal models of human disease, including reversing or reducing hepatic lipogenesis and steatosis in murine models of diet-induced obesity and non-alcoholic fatty liver disease.^1,2^. In macrophages, nitro-oleate reduced cholesterol accumulation by interacting with the CD36 scavenger receptor to reduce uptake of modified low density lipoproteins while increasing cholesterol efflux.^40,41^ Our study revealed that transcriptional regulators, lipid metabolism and transport proteins, including validated protein targets ESYT2 and RXRα, may be relevant cellular targets of nitro-fatty acids in living cells. ESYT2 has been implicated in the formation of ER-plasma membrane contact sites and lipid transfer between membranes.^42–44^ RXRα represents a central node in nuclear hormone receptor signaling and in the regulation of cellular metabolism by forming heterodimers with other nuclear receptors, including PPARs, liver X receptors and farnesoid X receptor.^45^ Furthermore, the presence of NCEH1, SOAT1 and SCARB1, which are key proteins in cholesterol hydrolysis, esterification and transport respectively, in our high-confidence list points to a possible mechanism by which nitro-fatty acids modulate cholesterol flux. Further investigations into how nitro-alkylation affects the function of these proteins may provide mechanistic insights into the biological effects of nitro-fatty acids in disease models characterized by dysfunctional cellular lipid metabolism such as atherosclerosis and metabolic syndrome.^46^

## 5. Conclusion

Here we demonstrated the stereo- and regio-selective synthesis of a clickable 9-nitro-oleate probe and successfully applied it to globally the profile protein targets of nitro-fatty acids in living macrophages. In addition to known targets, we identified novel protein targets of nitro-fatty acids that include important regulators of macrophage inflammatory responses as well as lipid metabolism and transport proteins. Overall, our study greatly expanded the scope of protein nitro-alkylation and provides a useful resource towards understanding the pleiotropic beneficial biological roles of nitro-fatty acids as signaling molecules and as multi-target therapeutic agents.

## Acknowledgements

We are grateful for financial support from NHRI and the Ministry of Science and Technology (M.M.Z.). The MS instrumental and data analysis resources were supported by the proteomic core lab (CLRPD1J0013 and 109-2113-M-182-003) at Chang Gung University, Taoyuan, Taiwan.

## Figures and Figure Legends

**Scheme 1.**
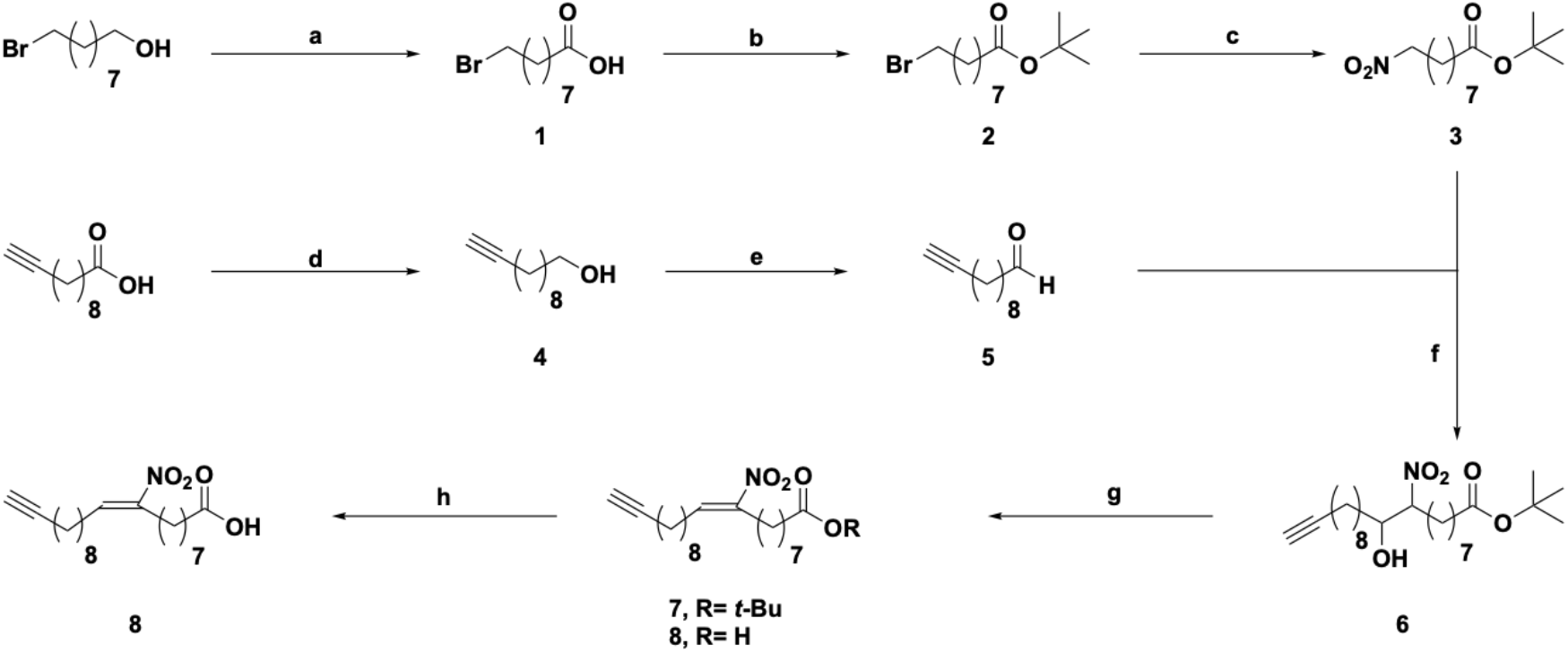
Synthesis of the clickable 9-nitro-oleate probe.

## References

1. Khoo, N. K. H. et al. Electrophilic nitro-oleic acid reverses obesity-induced hepatic steatosis. Redox Biology 22, 101132 (2019).

2. Rom, O. et al. Nitro-fatty acids protect against steatosis and fibrosis during development of nonalcoholic fatty liver disease in mice. EBioMedicine 41, 62–72 (2019).

3. Turell, L., Steglich, M. & Alvarez, B. The chemical foundations of nitroalkene fatty acid signaling through addition reactions with thiols. Nitric Oxide 78, 161–169 (2018).

4. Schopfer, F. J., Vitturi, D. A., Jorkasky, D. K. & Freeman, B. A. Nitro-fatty acids: New drug candidates for chronic inflammatory and fibrotic diseases. Nitric Oxide 79, 31–37 (2018).

5. Baker, L. M. S. et al. Nitro-fatty acid reaction with glutathione and cysteine. Kinetic analysis of thiol alkylation by a Michael addition reaction. J Biol Chem 282, 31085–31093 (2007).

6. Melo, T., Montero-Bullón, J.-F., Domingues, P. & Domingues, M. R. Discovery of bioactive nitrated lipids and nitro-lipid-protein adducts using mass spectrometry-based approaches. Redox Biology 23, 101106 (2019).

7. Lamas Bervejillo, M. et al. A FABP4-PPARγ signaling axis regulates human monocyte responses to electrophilic fatty acid nitroalkenes. Redox Biology 29, 101376 (2020).

8. Hansen, A. L. et al. Nitro-fatty acids are formed in response to virus infection and are potent inhibitors of STING palmitoylation and signaling. PNAS 115, E7768–E7775 (2018).

9. Charron, G. et al. Robust fluorescent detection of protein fatty-acylation with chemical reporters. J. Am. Chem. Soc. 131, 4967–4975 (2009).

10. Huang, D. W., Sherman, B. T. & Lempicki, R. A. Bioinformatics enrichment tools: paths toward the comprehensive functional analysis of large gene lists. Nucleic Acids Res 37, 1–13 (2009).

11. Huang, D. W., Sherman, B. T. & Lempicki, R. A. Systematic and integrative analysis of large gene lists using DAVID bioinformatics resources. Nat Protoc 4, 44–57 (2009).

12. Zhou, Y. et al. Metascape provides a biologist-oriented resource for the analysis of systems-level datasets. Nat Commun 10, 1523 (2019).

13. Morpheus. https://software.broadinstitute.org/morpheus. (Broad Institute).

14. Kauppi, B. et al. The three-dimensional structures of antagonistic and agonistic forms of the glucocorticoid receptor ligand-binding domain: RU-486 induces a transconformation that leads to active antagonism. J Biol Chem 278, 22748–22754 (2003).

15. Venkatachalam, C. M., Jiang, X., Oldfield, T. & Waldman, M. LigandFit: a novel method for the shape-directed rapid docking of ligands to protein active sites. J Mol Graph Model 21, 289–307 (2003).

16. Brooks, B. R. et al. CHARMM: the biomolecular simulation program. J Comput Chem 30, 1545–1614 (2009).

17. Bianco, G., Forli, S., Goodsell, D. S. & Olson, A. J. Covalent docking using autodock: two-point attractor and flexible side chain methods. Protein Sci 25, 295–301 (2016).

18. Parker, C. G. & Pratt, M. R. Click chemistry in proteomic investigations. Cell 180, 605–632 (2020).

19. Tsukidate, T., Li, Q. & Hang, H. C. Targeted and proteome-wide analysis of metabolite-protein interactions. Current Opinion in Chemical Biology 54, 19–27 (2020).

20. Khoo, N. K. H., Li, L., Salvatore, S. R., Schopfer, F. J. & Freeman, B. A. Electrophilic fatty acid nitroalkenes regulate Nrf2 and NF-κB signaling:A medicinal chemistry investigation of structure-function relationships. Scientific Reports 8, 2295 (2018).

21. Fazzari, M. et al. Electrophilic fatty acid nitroalkenes are systemically transported and distributed upon esterification to complex lipids. Journal of Lipid Research 60, 388–399 (2019).

22. Woodcock, S. R., Marwitz, A. J. V., Bruno, P. & Branchaud, B. P. Synthesis of nitrolipids. All four possible diastereomers of nitrooleic acids: (E)- and (Z)-, 9- and 10-nitro-octadec-9-enoic acids. Org. Lett. 8, 3931–3934 (2006).

23. Dunny, E. & Evans, P. Stereocontrolled synthesis of the PPAR-γ agonist 10-nitrolinoleic acid. J. Org. Chem. 75, 5334–5336 (2010).

24. Woodcock, S. R., Bonacci, G., Gelhaus, S. L. & Schopfer, F. J. Nitrated fatty acids: synthesis and measurement. Free Radic Biol Med 59, 14–26 (2013).

25. Hock, K. J. et al. Modular regiospecific synthesis of nitrated fatty acids. Synthesis 49, 615–636 (2017).

26. Kansanen, E. et al. Electrophilic nitro-fatty acids activate NRF2 by a KEAP1 cysteine 151-independent mechanism. J. Biol. Chem. 286, 14019–14027 (2011).

27. Sun, W. et al. ERIS, an endoplasmic reticulum IFN stimulator, activates innate immune signaling through dimerization. PNAS 106, 8653–8658 (2009).

28. Wang, X. et al. STING requires the adaptor TRIF to trigger innate immune responses to microbial infection. Cell Host Microbe 20, 329–341 (2016).

29. Woodcock, C.-S. C. et al. Nitro-fatty acid inhibition of triple-negative breast cancer cell viability, migration, invasion, and tumor growth. J Biol Chem 293, 1120–1137 (2018).

30. Davda, D. et al. Profiling targets of the irreversible palmitoylation inhibitor 2-bromopalmitate. ACS Chem. Biol. 8, 1912–1917 (2013).

31. Zheng, B. et al. 2-Bromopalmitate analogues as activity-based probes to explore palmitoyl acyltransferases. J. Am. Chem. Soc. 135, 7082–7085 (2013).

32. Weerapana, E. et al. Quantitative reactivity profiling predicts functional cysteines in proteomes. Nature 468, 790–795 (2010).

33. Abo, M. & Weerapana, E. A caged electrophilic probe for global analysis of cysteine reactivity in living cells. J. Am. Chem. Soc. 137, 7087–7090 (2015).

34. Salvatore, S. R., Vitturi, D. A., Fazzari, M., Jorkasky, D. K. & Schopfer, F. J. Evaluation of 10-nitro oleic acid bio-elimination in rats and humans. Sci Rep 7, (2017).

35. Fazzari, M. et al. Nitro-fatty acid pharmacokinetics in the adipose tissue compartment. J Lipid Res 58, 375–385 (2017).

36. Kansanen, E. et al. Nrf2-dependent and -independent responses to nitro-fatty acids in human endothelial cells. J Biol Chem 284, 33233–33241 (2009).

37. Schopfer, F. J. et al. Covalent peroxisome proliferator-activated receptor γ adduction by nitro-fatty acids: selective ligand activity and anti-diabetic signaling actions. J. Biol. Chem. 285, 12321–12333 (2010).

38. Ambrozova, G. et al. Nitro-oleic acid modulates classical and regulatory activation of macrophages and their involvement in pro-fibrotic responses. Free Radic Biol Med 90, 252–260 (2016).

39. Wang, P. et al. Electrophilic nitro-fatty acids suppress psoriasiform dermatitis: STAT3 inhibition as a contributory mechanism. Redox Biology 43, 101987 (2021).

40. Vazquez, M. M. et al. Nitro-oleic acid, a ligand of CD36, reduces cholesterol accumulation by modulating oxidized-LDL uptake and cholesterol efflux in RAW264.7 macrophages. Redox Biology 36, 101591 (2020).

41. Rosenblat, M., Rom, O., Volkova, N. & Aviram, M. Nitro-oleic acid reduces J774A.1 macrophage oxidative status and triglyceride mass: involvement of Paraoxonase2 and triglyceride metabolizing enzymes. Lipids 51, 941–953 (2016).

42. Saheki, Y. et al. Control of plasma membrane lipid homeostasis by the extended synaptotagmins. Nat Cell Biol 18, 504–515 (2016).

43. Yu, H. et al. Extended synaptotagmins are Ca2+-dependent lipid transfer proteins at membrane contact sites. Proc Natl Acad Sci U S A 113, 4362–4367 (2016).

44. Schauder, C. M. et al. Structure of a lipid-bound extended synaptotagmin indicates a role in lipid transfer. Nature 510, 552–555 (2014).

45. Evans, R. M. & Mangelsdorf, D. J. Nuclear receptors, RXR, and the big bang. Cell 157, 255–266 (2014).

46. Rom, O., Liu, Y., Chang, L., Chen, Y. E. & Aviram, M. Nitro fatty acids: novel drug candidates for the co-treatment of atherosclerosis and non-alcoholic fatty liver disease. Curr Opin Lipidol 31, 104–107 (2020).

47. Yang, J., Tallman, K. A., Porter, N. A. & Liebler, D. C. Quantitative chemoproteomics for site-specific analysis of protein alkylation by 4-hydroxy-2-nonenal in Cells. Anal. Chem. 87, 2535–2541 (2015).

48. Codreanu, S. G. et al. Alkylation damage by lipid electrophiles targets functional protein systems. Mol Cell Proteomics 13, 849–859 (2014).

