## Supporting Information for "Chemoproteomic profiling reveals cellular targets of nitro-fatty acids"

**Supplementary Figures** **Page S2-S3**

**Supplementary Tables Legends** **Page S4**

**Supplementary Materials and Methods** **Page S5-S11**

**Supplementary References** **Page S12**

### Supplementary Figures

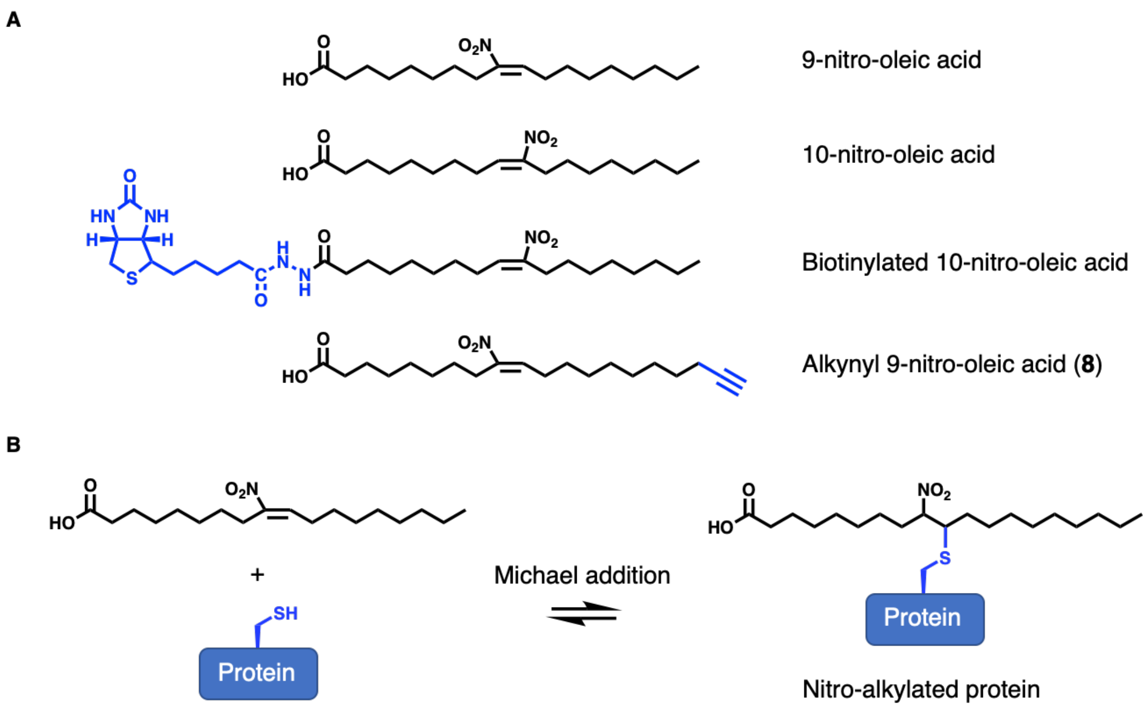


**Supplementary Figure 1. Protein nitro-alkylation is a reversible protein modification. (A)** Chemical structures of selected positional isomers of nitro-oleic acids, 9-nitro-oleate and 10-nitro-oleate. Also shown are the chemical structures of nitro-oleate probes for the detection of nitro-alkylated proteins, including the widely used biotinylated nitro-oleate,[1] and the alkynyl-9-nitro-oleate probe **8** from this study. **(B)** Nitro-fatty acids form reversible protein adducts with the thiols of cysteine residues of target proteins via Michael addition.

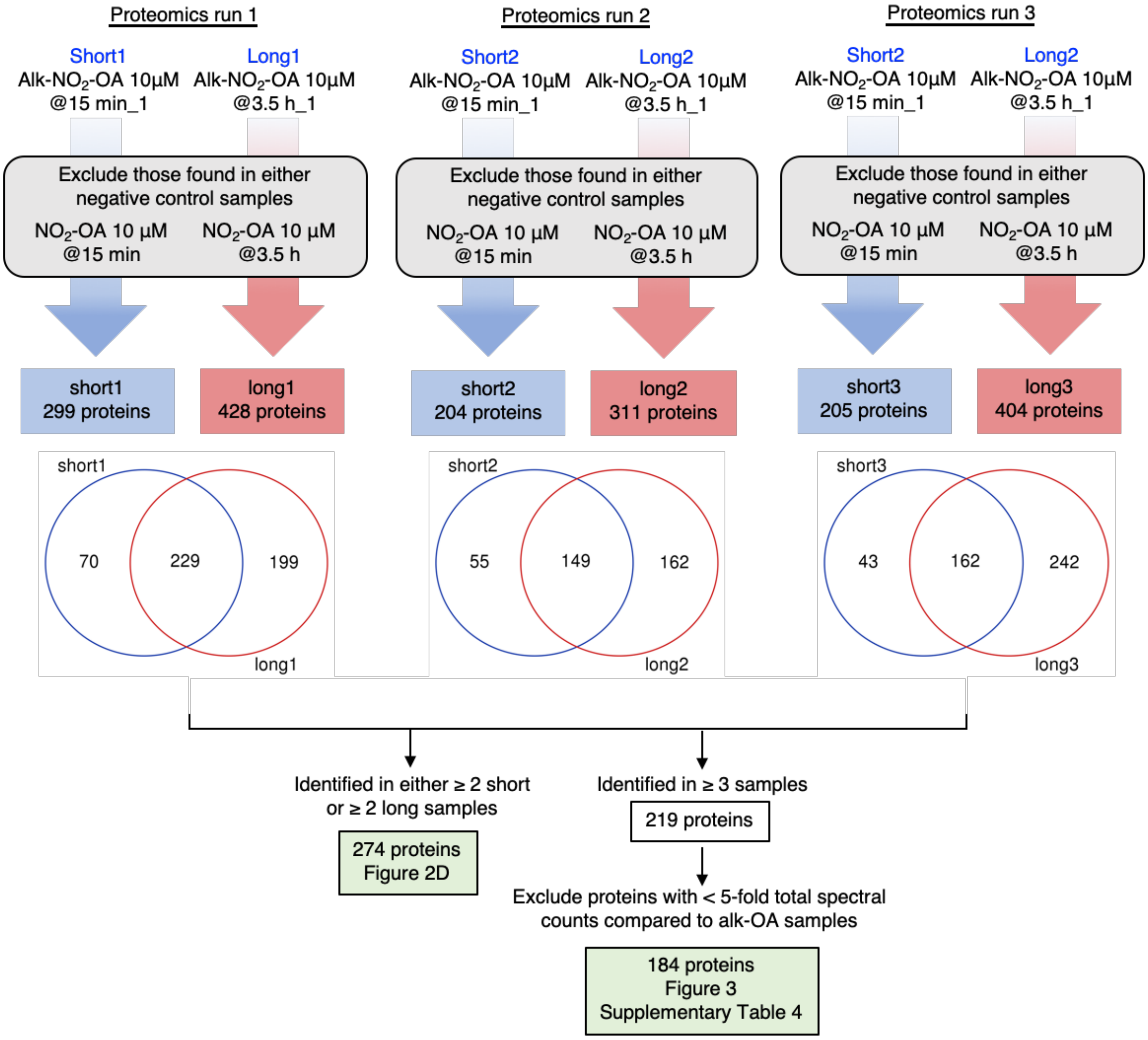


**Supplementary Figure 2. Filter criteria for the identification of nitro-alkylated proteins from THP1 cells.** For each proteomics run, from the list of proteins obtained in the alk-9-$NO_2$-OA samples that were treated for 15 min (short) and 3.5 h (long), we excluded proteins that were also identified in either of the two negative control samples (9-$NO_2$-OA). Filtering for proteins that were either identified in $\geq 2$ short or $\geq 2$ long samples yielded 274 proteins as represented in Figure 2D. Filtering for proteins that were identified $\geq 3$ times in the total of 6 samples yielded and excluding proteins that were similarly enriched using alk-OA yielded 184 proteins. These proteins were used for bioinformatics analyses represented in Figure 3 and listed in Supplementary Table 1. For more description of the filter criteria, refer to Methods.

### Supplementary Tables Legends

**Supplementary Table 1. High-confidence list of nitro-alkylated proteins from THP1 macrophages identified by chemoproteomics study.** Proteins are ranked by the total number of PSMs identified in alk-9-$NO_2$-OA samples. (.xlxs)

**Supplementary Table 2. Comparative anaylsis of high confidence nitro-alkylated protein list from this study with protein targets of other indicated electrophilic compounds.** Proteins found to be modified by the indicated electrophiles in their respective studies are highlighted in green. IA, iodoacetamide.[2] BMK, bromomethyl ketone.[3] HNE, 4-hydroxy-2-nonenal and ONE, 4-oxo-2-nonenal.[4,5] 2BP, 2-bromopalmitate.[6,7] (.xlsx)

**Supplementary Materials and Methods**

**Chemical synthesis of clickable alk-9-$NO_2$-OA probe.**

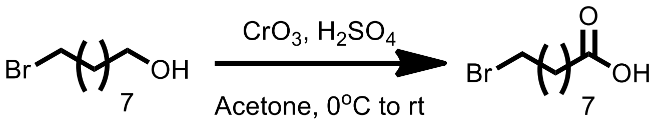


**1**

**9-bromononanoic acid 1**: To the solution of 2.5 M Jones' reagent (30 mL, 76.18 mmol, 1.7 eq.) in 10 mL of acetone was added 9-bromononanol (10 g, 44.81 mmol) in acetone (10 mL) at 0 °C. After reaction was completed, the solution mixture was added water (100 mL) and $Et_2O$ (100 mL). After phase separation, the aqueous layer was extracted with $Et_2O$ (50 mL x 3). The combined organic layer was washed with 1.0 N NaOH (30 mL x 3). The aqueous layers were combined and washed with DCM (30 mL x 2) to remove nonpolar impurities. The aqueous layer was acidified to pH 2 with conc. HCl and extracted with $Et_2O$ (30 mL x 3). The combined organic layers were washed with brine (50 mL x 2), dried over $MgSO_4$, filtered, and concentrated in vacuo to provide 9-bromononanoic acid **1** (6.68 g, 63%). $^{1}$H NMR ($CDCl_3$, 400 MHz): δ 3.38 (2H, t, $J$ = 9.2 Hz), 2.33 (2H, $J$ = 10 Hz), 1.88-1.78 (2H, m), 1.64-1.57 (2H, m), 1.42-1.40 (2H, m), 1.38-1.30 (6H, m); $^{13}$C NMR ($CDCl_3$, 176 MHz): δ 180.51, 34.20, 34.04, 32.88, 29.14, 29.02, 28.65, 28.19, 24.71.[8]
Jone's reagent preparation: Dissolve $CrO_3$ (25 g, 0.25 mol) in $H_2O$ (75 mL) then was added conc. $H_2SO_4$ (25 mL) slowly at 0 °C. The concentration of the solution prepared by this procedure is 2.5 M.

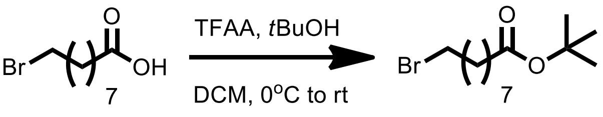


**1** **2**

**Ester 2:** To a solution of 9-bromononanoic acid **1** (208 mg, 0.88 mmol) in DCM (4.4 mL) was added dropwise TFAA (405 mg, 1.93 mmol, 2.2 eq.) at 0 °C. The resulting mixture was stirred for 2.5 h before addition *t*BuOH (228 mg, 3.07 mmol, 3.5 eq.) at 0 °C. The resulting mixture was slowly warmed to RT. After reaction was completed, resulting mixture was added 10 mL of $NaHCO_3$ (sat. aq) and extracted with $Et_2O$ (10 mL x 3). The combined organic layers were washed with brine, dried over $MgSO_4$, filtered, and concentrated in vacuo to give ester **2** (238 mg, 93%). $^{1}$H NMR (400 MHz, $CDCl_3$): δ 3.38 (2H, t, $J$ = 7.2 Hz), 2.18 (2H, 5, $J$ = 7.6 Hz), 1.86-1.79 (2H, m), 1.59-1.52 (overlap with water residual, 2H, m), 1.43-1.38 (11H, m), 1.29 (6H, br); $^{13}$C NMR (176 MHz, $CDCl_3$): δ 173.20, 79.91, 35.59, 33.93, 32.85, 29.16, 29.02, 28.65, 28.19, 28.17, 25.09.[9]

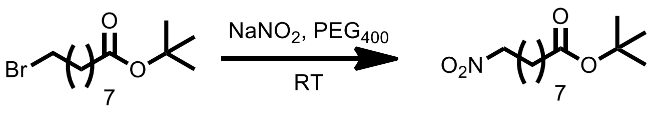


**2** **3**

***tert*-butyl 9-nitrononanoate 3**: To the flask was added $NaNO_2$ (424 mg, 6.15 mmol, 3.0 eq.) and PEG-400 (4.0 mL). The resulting mixture was stirred for 3 h at RT before addition ester **2**

(600 mg, 2.05 mmol). After the reaction was completed, resulting mixture was added 40 mL of water and extracted with $Et_2O$ (20 mL x 3). The combined organic layers were washed with brine, dried over $MgSO_4$, filtered, and concentrated in vacuo. The residue was purified by flash chromatography (10% EtOAc/hexane, $R_f$=0.33) to yield the *tert*-butyl 9-nitrononanoate **3** (234 mg, 44%). $^1$H NMR (400 MHz, $CDCl_3$): δ 4.351 (2H, t, *J* = 7.2Hz), 2.18 (2H, t, *J* = 7.6), 1.99-1.94 (2H, m), 1.58-1.52(2H, m), 1.42 (9H, s), 1.37-1.29 (8H, m); $^{13}$C NMR (176 MHz, $CDCl_3$): δ 172.96, 79.72, 75.47, 35.28, 28.73, 28.66, 28.49, 27.90, 27.16, 25.96, 24.77; HRMS (ESI): calculated for $C_{13}H_{25}NNaO_4$ [M+Na$^+$] 282.1676, found 282.1690.

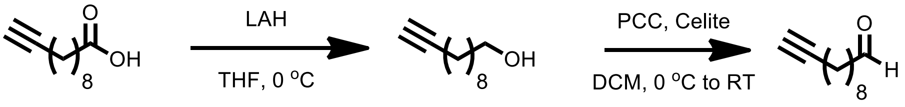


**4** **5**

To a solution of LAH (603 mg, 15.09 mmol) in THF (40 mL) was added dropwise undec-10-ynoic acid (1338 mg, 7.95 mmol) in THF (10 mL) at 0 °C. After the reaction was completed, resulting mixture was added 10 mL of water at 0 °C and then adjusted to pH 3-5 using conc. HCl. The aqueous layer was extracted with $Et_2O$ (100 mL x 3). The combined organic layers were washed with brine, dried over $MgSO_4$, filtered, and concentrated in vacuo to give crude alcohol **4**. The crude alcohol **4** was used for next reaction without further purification. $^1$H NMR (400 MHz, $CDCl_3$): δ 3.64 (t, J = 6.6 Hz, 2H), 3.49 (s, 1H), 2.18 (dt, J = 7.1, 2.6 Hz, 2H), 1.94 (t, J = 2.7 Hz, 1H), 1.63 – 1.47 (m, 4H), 1.46 – 1.21 (m, 10H); $^{13}$C NMR (100 MHz, $CDCl_3$): δ 84.92, 68.20, 63.20, 32.91, 29.57, 29.49, 29.16, 28.85, 28.60, 25.84, 18.53.

Alcohol **4** was dissolved in DCM (40 mL) followed by addition of PCC (2.23 g, 10.34 mmol) and celite (2.6 g) at 0 °C. After the reaction was completed, resulting mixture was filtered by celite and concentrated in vacuo. The residue was purified by flash chromatography (10%, EtOAc/hexane, $R_f$=0.53) to yield the aldehyde **5** (759 mg, 57% in two steps).[10] $^1$H NMR (400 MHz, $CDCl_3$): δ 9.74 (1H, t, *J* = 1.6Hz), 2.40 (2H, td, *J* = 7.36, 1.84Hz), 2.16 (2H, td, *J* = 7.0, 2.64Hz), 1.92 (1H, t, *J* = 2.64Hz), 1.64-1.59 (3H, m), 1.53-1.46(2H, m), 1.40-1.35 (2H, m), 1.33-1.26(5H, m); $^{13}$C NMR (100 MHz, $CDCl_3$): δ 203.163, 84.901, 68.324, 44.092, 29.407, 29.285, 29.071, 28.842, 28.621, 22.247, 18.579.[10]

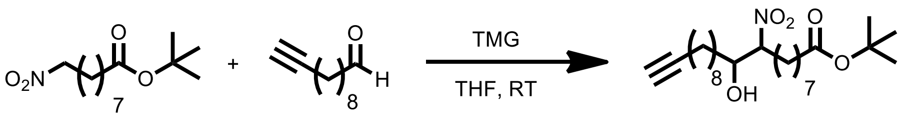


**3** **5** **6**

**β-Hydroxynitro ester 6:** To a solution of *tert*-butyl 9-nitrononanoate **3** (299 mg, 1.15 mmol) and aldehyde **5** (211 mg, 1.27 mmol, 1.1 eq.) in THF (6.0 mL) was added TMG (27 mg, 0.23 mmol, 0.2 eq.) dropwise at RT. After the reaction was completed, solvent was removed by reducing pressure and the residue was purified by flash chromatography (10% EtOAc/hexane, $R_f$=0.08) to yield the 45:55 mixture of diastereomers β-hydroxynitro ester **6** (332 mg, 68%). The mixture was used for next reaction without further purification.

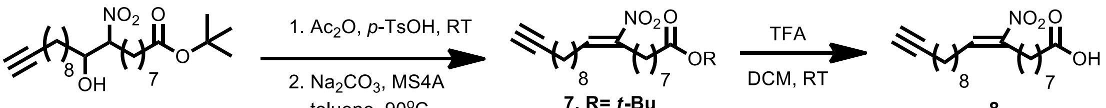


**6** **7, R= *t*-Bu** **8, R= H** **8**

**((*E*)-9-nitroicos-9-en-19-ynoic acid 8:** The β-hydroxynitro ester **6** (205 mg, 0.48 mmol) in round bottom flask was added acetic anhydride (3.0 mL) as solvent and a catalytic amount of *p*-toluenesulfonic acid (10 mg). The solution was stirred under $N_2$ at RT for overnight. After reaction was completed, the solvent was removed by azeotropic removal with toluene (3 × 20 mL) under reduced pressure. The residual was dissolved in toluene (3 mL) and addition of $Na_2CO_3$ (51 mg, 0.482 mmol, 1.0 eq.). The solution was heated to 90 °C under $N_2$ for overnight. After the reaction was completed, the solution is cooled to RT and diluted with 10 mL of $NH_4Cl$ (sat.aq) and 10 mL $Et_2O$. After phase separation, the aqueous layer is extracted with $Et_2O$ (10 mL x 3). The combined organic layers were washed with brine, dried over $MgSO_4$, filtered, and concentrated in vacuo to afford nitroalkene products **7** and **8** as a mixture. The mixture was used for next reaction without further purification.

To the above mixture in DCM (3.0 mL) was added TFA (1.0 mL) at RT. After the reaction was completed, the solvent and excess TFA were removed by reducing pressure. The residual was purified by flash chromatography (20% EtOAc/hexane, $R_f$=0.42) to yield the (*E*)-9-nitroicos-9-en-19-ynoic acid **8** (69 mg, 41% in three steps). $^1H$ NMR (400 MHz, $CDCl_3$): δ 7.08 (1H, t, *J* = 7.6 Hz), 2.56 (2H, appear t), 2.35 (2H, t, J = 7.48), 2.24-2.16 (4H, m), 1.94 (1H, t, J = 2.64), 1.65-1.61 (3H, m), 1.54-1.44 (8H, m), 1.41-1.25(11H, m); $^{13}C$ NMR (100 MHz, $CDCl_3$): δ 180.0, 151.7, 136.4, 84.6, 68.1, 33.9, 29.2, 29.1, 28.9, 28.84, 28.80, 28.6, 28.4, 28.3, 27.9, 27.8, 26.2, 24.5; HRMS (ESI): calculated for $C_{20}H_{32}NO_4$ [M−H$^+$] 350.2337, found 350.2321.

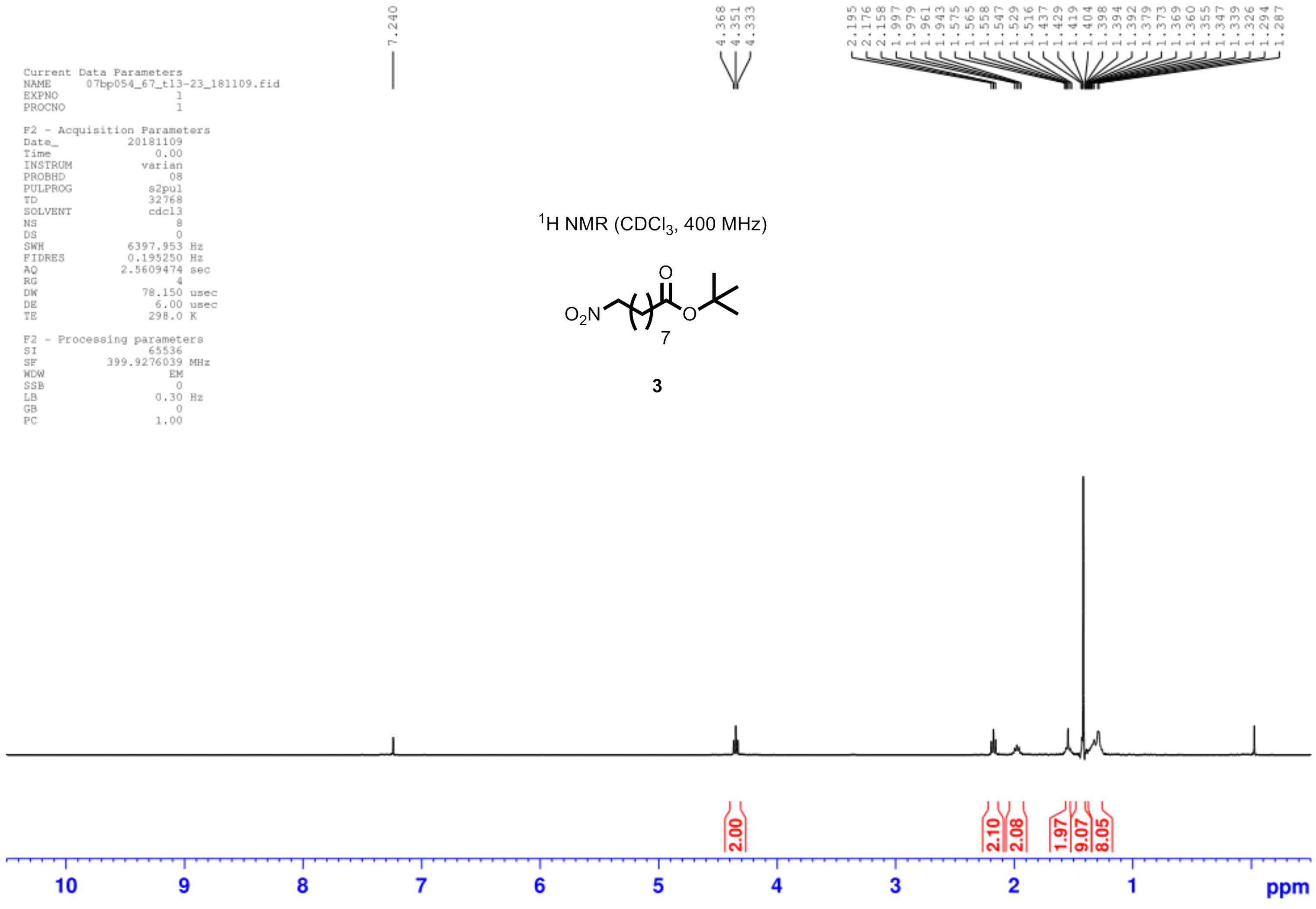
Current Data Parameters
NAME 07bp054_67_t13-23_181109.fid
EXPNO 1
PROCNO 1
F2 - Acquisition Parameters
Date_ 20181109
Time 0.00
INSTRUM varian
PROBHD 08
PULPROG s2pul
TD 32768
SOLVENT cdcl3
NS 8
DS 0
SWH 6397.953 Hz
FIDRES 0.195250 Hz
AQ 2.5609474 sec
RG 4
DW 78.150 usec
DE 6.00 usec
TE 298.0 K
F2 - Processing parameters
SI 65536
SF 399.9276039 MHz
WDW EM
SSB 0
LB 0.30 Hz
GB 0
PC 1.00
7.240
4.368
4.351
4.333
2.195
2.176
2.158
1.997
1.979
1.961
1.943
1.575
1.565
1.558
1.547
1.529
1.516
1.437
1.429
1.419
1.404
1.398
1.394
1.392
1.379
1.373
1.369
1.360
1.355
1.347
1.339
1.326
1.294
1.287
$^1$H NMR ($CDCl_3$, 400 MHz)
$O_2N$
O
O
7
3
2.00
2.10
2.08
1.97
9.07
8.05
10
9
8
7
6
5
4
3
2
1
ppm

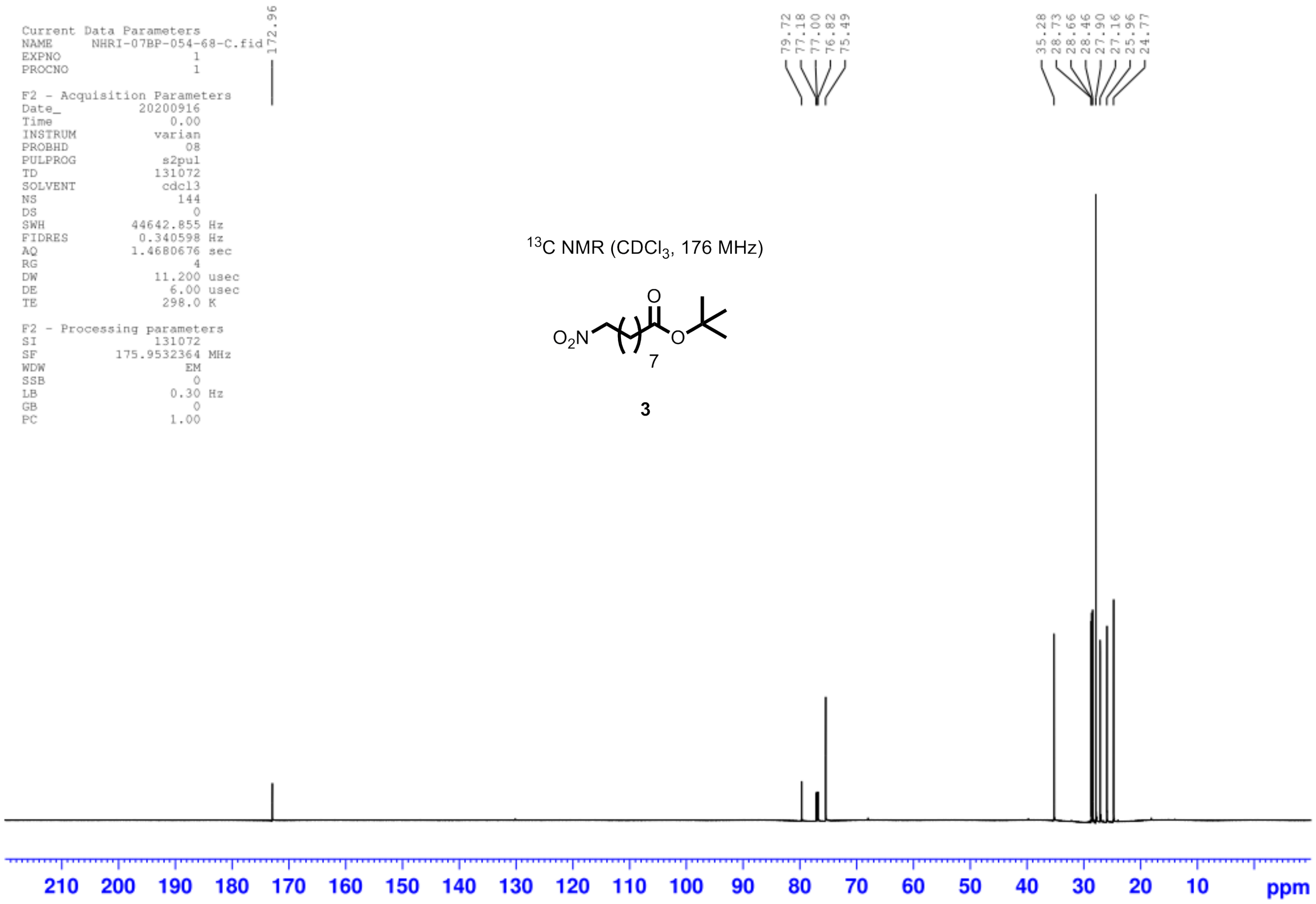

Current Data Parameters
NAME NHRI-07BP-054-68-C.fid
EXPNO 1
PROCNO 1
F2 - Acquisition Parameters
Date_ 20200916
Time 0.00
INSTRUM varian
PROBHD 08
PULPROG s2pul
TD 131072
SOLVENT cdcl3
NS 144
DS 0
SWH 44642.855 Hz
FIDRES 0.340598 Hz
AQ 1.4680676 sec
RG 4
DW 11.200 usec
DE 6.00 usec
TE 298.0 K
F2 - Processing parameters
SI 131072
SF 175.9532364 MHz
WDW EM
SSB 0
LB 0.30 Hz
GB 0
PC 1.00
172.96
79.72
77.18
77.00
76.82
75.49
35.28
28.73
28.66
28.46
27.90
27.16
25.96
24.77
13C NMR (CDCl3, 176 MHz)
O2N
O
O
7
3
210
200
190
180
170
160
150
140
130
120
110
100
90
80
70
60
50
40
30
20
10
ppm

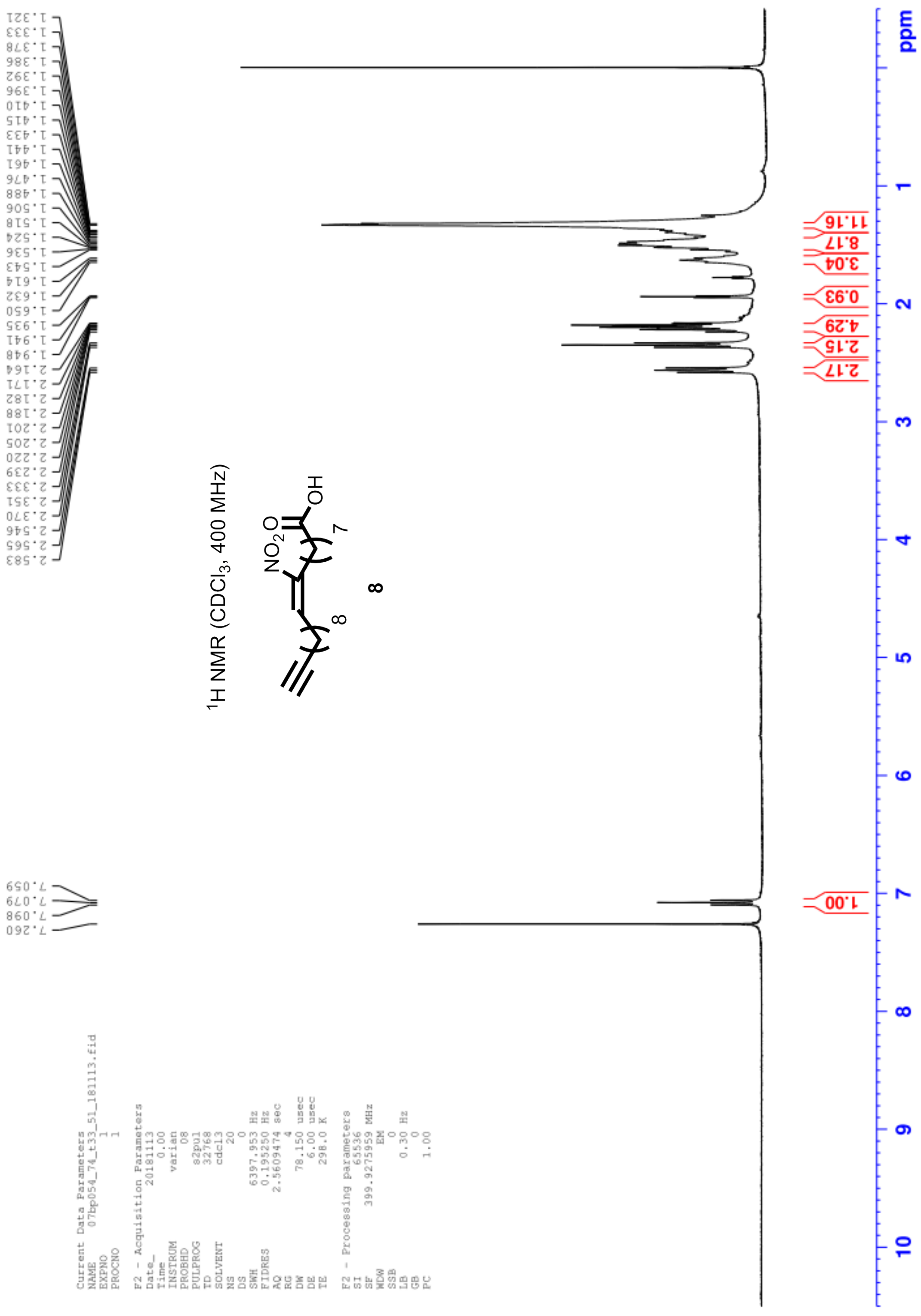
1H NMR (CDCl3, 400 MHz)
8
ppm

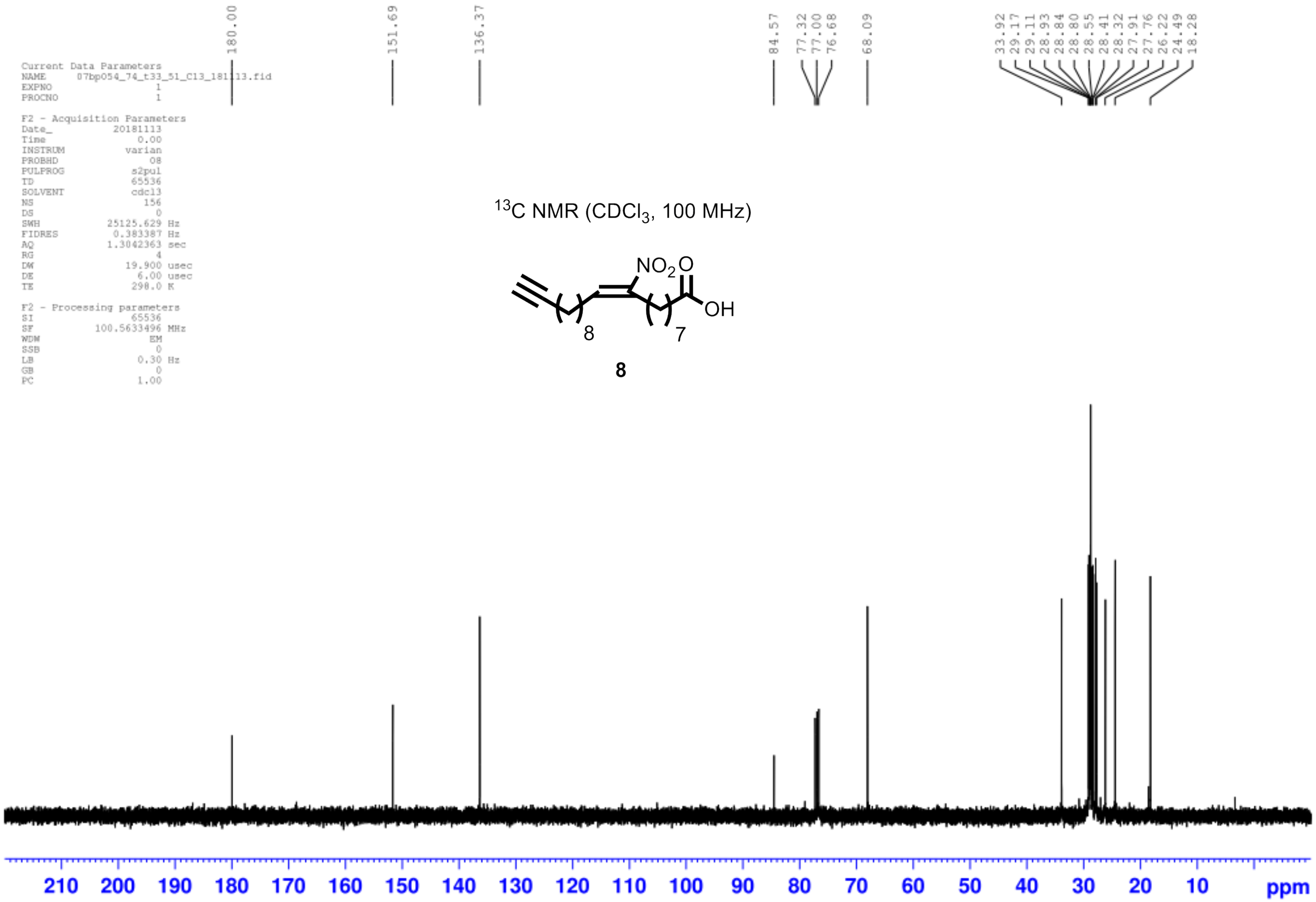
Current Data Parameters
NAME 07bp054_74_t33_51_C13_181113.fid
EXPNO 1
PROCNO 1
F2 - Acquisition Parameters
Date_ 20181113
Time 0.00
INSTRUM varian
PROBHD 08
PULPROG s2pul
TD 65536
SOLVENT cdcl3
NS 156
DS 0
SWH 25125.629 Hz
FIDRES 0.383387 Hz
AQ 1.3042363 sec
RG 4
DW 19.900 usec
DE 6.00 usec
TE 298.0 K
F2 - Processing parameters
SI 65536
SF 100.5633496 MHz
WDW EM
SSB 0
LB 0.30 Hz
GB 0
PC 1.00
180.00
151.69
136.37
84.57
77.32
77.00
76.68
68.09
33.92
29.17
29.11
28.93
28.84
28.80
28.55
28.41
28.32
27.91
27.76
26.22
24.49
18.28
$^{13}C$ NMR ($CDCl_3$, 100 MHz)
$NO_2$
O
OH
8
7
8
210 200 190 180 170 160 150 140 130 120 110 100 90 80 70 60 50 40 30 20 10 ppm

**References**


1. Cui, T. *et al.* Nitrated fatty acids: endogenous anti-inflammatory signaling mediators. *J. Biol. Chem.* **281**, 35686–35698 (2006).
2. Weerapana, E. *et al.* Quantitative reactivity profiling predicts functional cysteines in proteomes. *Nature* **468**, 790–795 (2010).
3. Abo, M. & Weerapana, E. A caged electrophilic probe for global analysis of cysteine reactivity in living cells. *J. Am. Chem. Soc.* **137**, 7087–7090 (2015).
4. Yang, J., Tallman, K. A., Porter, N. A. & Liebler, D. C. Quantitative chemoproteomics for site-specific analysis of protein alkylation by 4-hydroxy-2-nonenal in cells. *Anal. Chem.* **87**, 2535–2541 (2015).
5. Codreanu, S. G. *et al.* Alkylation damage by lipid electrophiles targets functional protein systems. *Mol Cell Proteomics* **13**, 849–859 (2014).
6. Davda, D. *et al.* Profiling targets of the irreversible palmitoylation inhibitor 2-bromopalmitate. *ACS Chem. Biol.* **8**, 1912–1917 (2013).
7. Zheng, B. *et al.* 2-Bromopalmitate analogues as activity-based probes to explore Palmitoyl acyltransferases. *J. Am. Chem. Soc.* **135**, 7082–7085 (2013).
8. Kumari, A., Gholap, S. P. & Fernandes, R. A. Tandem IBX-promoted primary alcohol oxidation/opening of intermediate β,γ-diolcarbonate aldehydes to (E)-γ-hydroxy-α,β-enals. *Chemistry – An Asian Journal* **14**, 2278–2290 (2019).
9. Baalmann, M. *et al.* A bioorthogonal click chemistry toolbox for targeted synthesis of branched and well-defined protein–protein conjugates. *Angewandte Chemie International Edition* **59**, 12885–12893 (2020).
10. Cheng, X., Li, L., Uttamchandani, M. & Yao, S. Q. In situ proteome profiling of C75, a covalent bioactive compound with potential anticancer activities. *Org. Lett.* **16**, 1414–1417 (2014).
